# Vertically Integrated System for Tracking and Assessing cell-cycle aware phenotypes under confinement

**DOI:** 10.1101/2025.09.25.678497

**Authors:** Melissa Pezzotti, Eloisa Torchia, Julius Zimmermann, Sara Rigolli, Alessandro Enrico, Moises Di Sante, Francesco S. Pasqualini

## Abstract

Quantitative cell biology often studies migration and the cell-cycle (CC) in separate assays, limiting mechanistic insights, particularly under geometric confinement. Here, we introduce a vertically integrated platform for simultaneously tracking single-cell migration and assessing CC under confinement. Our system integrates cell engineering via multiplexed sensors for cell-cycle, actin, and tubulin, as well as photopatterned engineered extracellular matrix (ECM) islands of defined sizes. It also features an automated, high-throughput pattern-aware imaging pipeline (Fab2Mic) that enables on-pattern, joint migration-CC assessment in the same live cells. Since the local microenvironment plays a critical role in metastasis by constraining cell behaviors within spatial boundaries, we used an HT1080 fibrosarcoma model as an illustrative case. Where static phenotyping yielded 40% G1 and 60% S/G2/M, with larger cell areas and tubulin spread in the S/G2/M phase, dynamic phenotyping via live-cell imaging confirmed CC-linked motility, with faster instantaneous velocities in G1, exemplifying the CC-migration correlations. These phenotypes were modulated by the spatial confinement imposed by the engineered ECM islands. Stronger confinement reduced cell area and tubulin spread and increased the frequency of abnormal CC events, particularly Long G1 states on smaller engineered ECM islands. It also induced a confinement-specific S/G2/M-G1 mitotic slippage, observed only under our confined conditions. Together, this vertically integrated system suggests that confinement may continuously tune migration–CC coupling and provides a deployable pipeline for CC-aware mechanobiology and screening. Moreover, we stress how dynamic imaging provides access to variables that are difficult or impossible to infer from static snapshots, including velocity and CC timing.

## Introduction

Recent estimates still report cancer as a leading cause of mortality, with 20 million new cases and 9.7 million deaths in 2022, the majority after metastatic spread^1^. Beyond genetic alteration, tumor progression is critically shaped by the tumor microenvironment, where the extracellular matrix (ECM), stromal barriers, and tissue architecture impose physical constraints on cell behaviors. In vivo, these microenvironmental features govern how cells migrate, invade, and proliferate during dissemination^2^. Among them, geometric confinement challenges cell migration^3^, affects nuclear integrity^4,5^, and can alter cell-cycle (CC) progression^6,7^. However, in vitro assays generally measure either migration or proliferation: standard 2D assays, such as scratch and transwell assays^8–10^, report collective migration or transmigration but do not detail cell-cycle phase; more advanced 3D in vitro models (e.g., spheroids, collagen invasion assays) better mimic tissue architecture but likewise seldom provide real-time information on cell-cycle stage^11^. Conversely, endpoint readouts such as EdU and Ki-67 provide snapshots of the proliferative state of the cells but cannot track CC dynamics over time^12,13^. Furthermore, ECM cues, such as substrate stiffness^13–15^, chemical composition with force transmission^16^, and cell shape and geometry^17^, have been shown to regulate adhesion, cytoskeletal tension^18–20^, and proliferation programs^21–24^. Each of these has been linked to either proliferative or migratory phenotypes, underscoring the importance of contextual mechanical inputs in determining cell fate.

To study how confinement reshapes CC-migration coupling^7,25–27^, it is crucial to integrate confined migration assays with live CC readouts. However, open technical challenges are preventing this integration. First, most fluorescent CC reporters rely on GFP and RFP^28^, the same spectral channels used by common structural and functional sensors. This competition limits multiplexing, making it difficult to track the CC and migration-driving cytoskeletal dynamics simultaneously. Second, photopatterning approaches such as LIMAP and PRIMO now allow fabrication of engineered ECM islands with precise geometry^29^. However, aligning these patterns with microscope fields of view still requires manual trial-and-error, reducing throughput and reproducibility^30^. Finally, it has been difficult to obtain accurate long-term quantification when cells can move out of the field-of-view over hours of imaging^31–33^. These obstacles span distinct competencies: reporter engineering, photopatterned microfabrication, and high-content imaging and analysis, which require different skill sets and infrastructure. Bringing them together within one workflow imposes a substantial coordination and tooling burden, which has constrained routine, large-throughput execution of end-to-end CC–aware confinement assays.

To address this gap, we introduce a vertically integrated platform that simultaneously tracks migration and assesses the cell-cycle, unifying multiplexed reporters, photopatterned microfabrication, and high-content live-cell imaging. This system builds directly on our CALIPERS^34^ framework, which leverages FUCCIplex, a cell-cycle sensor spectrally multiplexable with diverse structural and functional reporters. Here, we extend this concept to a migratory HT1080 fibrosarcoma model and integrate it with Fab2Mic. This fabrication-to-microscopy correlative pipeline engineers ECM islands and registers them to acquisition coordinates, enabling automated imaging of single-cell migration and proliferation on each island. Custom image-analysis scripts segment and track individual cells and nuclei over time, allowing quantification of how geometric confinement modulates migration–CC coupling.

## Results

### Microscopy-based automatic selection for engineered reporter line

We set out to engineer HT1080 fibrosarcoma cells into a four-reporter reference line that simultaneously encodes CC state and migration-supporting cytoskeletal structure (Fig. 1A). In this work, we used HT1080 cells from Ibidi that already carried a green fluorescent protein (EGFP) LifeAct construct. To complement our four-color reporter line, we edited the endogenous TUBA1B locus with a red fluorescent protein (RFP) tag via CRISPR/Cas9 editing and introduced the FUCCIplex cassette via lentiviral infection under the EF1α promoter (see cell line generation in the Methods section). The FUCCIplex design replaces the RFP and GFP markers of G1 and S/G2/M phases in traditional FUCCI sensors^28^ with cyan and far-red variants (CFP and iRFP), respectively.

**Figure 1.**
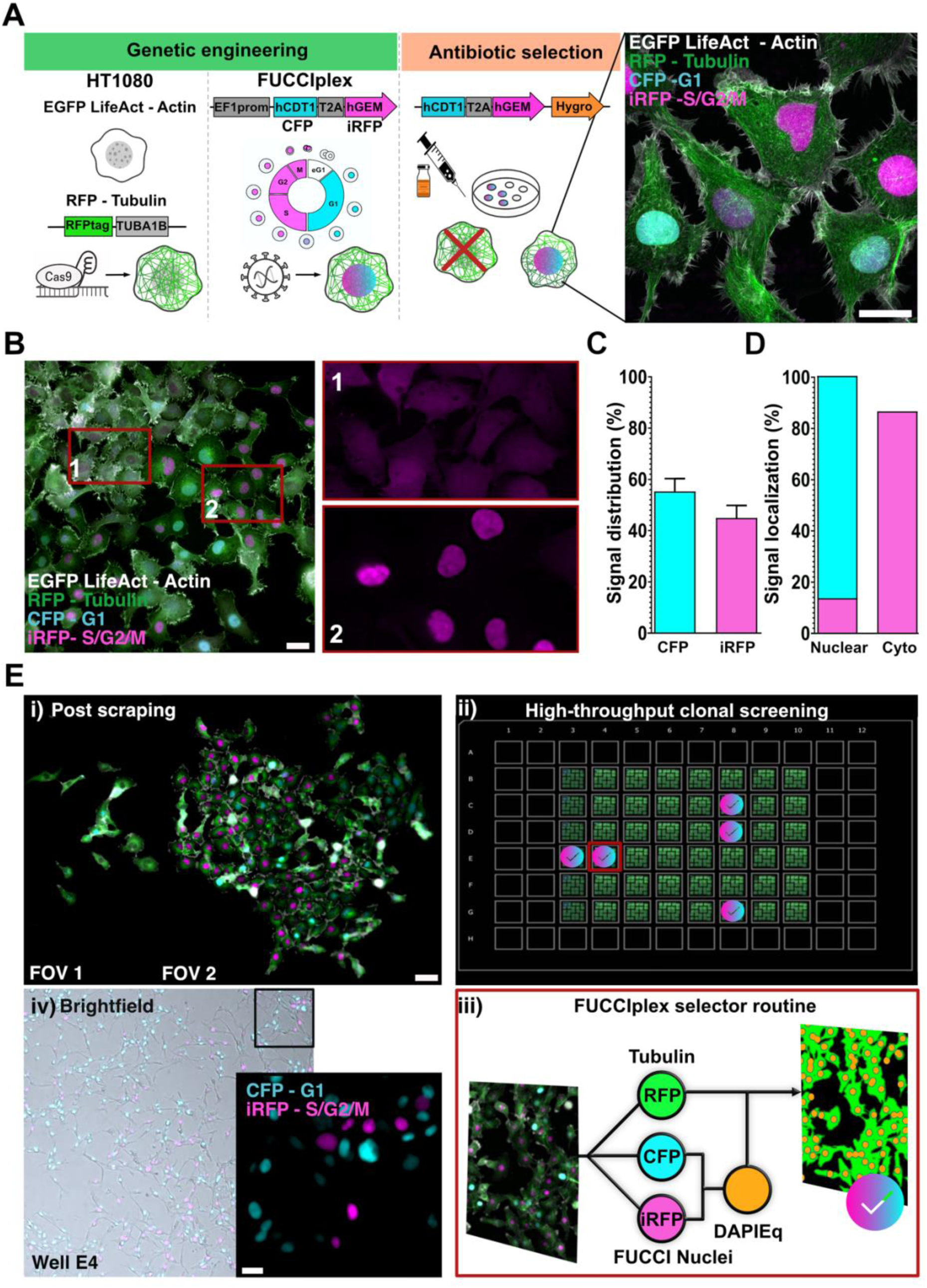
Engineering and rescue of a four-reporter CALIPERS-enabled HT1080 reference line. (A) Schematic of the genetic engineering pipeline. HT1080 fibrosarcoma cells pre-engineered with EGFP LifeAct were genome-edited at the endogenous TUBA1B locus to introduce RFP-tubulin and transduced with the FUCCIplex cassette (CFP-G1 and iRFP-S/G2/M). Lentiviral infection and hygromycin selection yielded a mixed population expressing four channels (actin, tubulin, G1, S/G2/M). High-resolution confocal imaging confirmed robust actin filaments, tubulin networks, and phase-marked nuclei. (B) Widefield imaging revealed anomalous cytoplasmic iRFP signal in many cells (example in inset 1) compared to correctly localized nuclear iRFP (inset 2). (C) Quantification of signal distribution in the mixed population (n = 898 cells, 26 FOVs) showed 55% ± 5% of total cells CFP+ and 45% ± 5% of total cells iRFP+. (D) Localization analysis confirmed 100% nuclear localization of CFP, whereas only 15% of the iRFP signal was nuclear, with the majority retained in the cytoplasm. (E) Rescue pipeline. (i) Post-scraping enrichment of cells with nuclear iRFP yielded approximately 3× higher nuclear signal. (ii) Clonal seeding of the scraped population in 96-well format enabled systematic high-throughput screening. (iii) An automated selector routine segmented tubulin as ground truth and generated a DAPI-equivalent mask from CFP and iRFP nuclear signals, calling a clone only when nuclear counts matched > 80% of the tubulin-defined cell count. (iv) Manual inspection of brightfield and fluorescence confirmed nuclear expression of both CFP and iRFP in selected clones. Five reference lines were banked, with clone E4 chosen as the working HT1080 CALIPERS line.

Following the selection of engineered cells using hygromycin, we confirmed that actin filaments, microtubules, and nuclei, marked in G1 (CFP) or S/G2/M (iRFP), were fluorescently tagged using high-resolution microscopy and PCR (Fig. 1A, Fig. S1A, C). In terms of nuclear markers, the mixed population featured CFP localized robustly to nuclei. Still, unexpectedly, iRFP was frequently retained in the cytoplasm (Fig. 1B). In cancer cells, such as HT1080, nuclear localization failures can arise from multiple causes, ranging from altered nuclear import machinery^4,5,35^ to deregulated protein degradation^36^. To assess the cell phenotype distribution, we annotated 898 cells across 26 fields of view. On average, 55% ± 5% of the total cell population expressed CFP and 45% ± 5% expressed iRFP (Fig. 1C). While all CFP signals localized to nuclei, only 15% ± 5% of iRFP+ cells exhibited a signal confined to the nucleus, while the remainder retained signal in the cytoplasm (Fig. 1D).

To enrich the engineered cell population for proper localization of iRFP, we mechanically scraped regions featuring cells with nuclear iRFP signal, expanded the population, and observed a roughly 3-fold increase in the frequency of cells with nuclear iRFP signal (Fig. 1E-i). We then clonally seeded the enriched population in 96-well plates and screened 40 resulting clones with a custom-made high-throughput acquisition via a Nikon JOB routine (Fig. 1E-ii). The automated selector routine works by segmenting tubulin to identify cells and then building a DAPI-equivalent mask from CFP and iRFP nuclear signals and calls an HT1080 CALIPERS four-reporter clone only when nuclear counts match > 80% of the tubulin count in the same field of view (Fig. 1E-iii). More than half the clones passed this filter, confirming the efficacy of the rescue strategy. Manual inspection of brightfield and fluorescence overlays validated the algorithm’s calls (Fig. 1E-iv). Ultimately, we banked five four-color reporter lines and designated clone E4 as the working HT1080 CALIPERS line (Fig. S1D).

### Baseline simultaneous assessment of cell-cycle (CC) progression and migration

To establish a reference baseline for CC-aware migratory phenotyping in free space, we first imaged HT1080 CALIPERS cells seeded on fibronectin-coated glass substrates. High-resolution fluorescence imaging confirmed all four colors could be detected, clearly highlighting the cell nucleus, with G1 cells marked by CFP and S/G2/M cells marked by iRFP, alongside actin and tubulin structure (Fig. 2A). Qualitatively, our analysis confirmed how cellular heterogeneity arises directly from the combined effects of cell-cycle state and morphology. Compared to Cell 1 (G1), Cell 2 (S/G2/M) had a considerably larger projected area. Conversely, Cell 3 shared the same S/G2/M phase as Cell 2, but it had a considerably smaller area, underscoring the variability present even within the same phase. We then quantified CC distribution and morphology across three independent experiments at early (4 h) and late (24 h) time points to control for seeding bias. A total of 60 cells were analyzed in the early condition and 82 in the late condition. G1 and S/G2/M comprised, respectively, approximately 37–39% and 58–61% of the population at both time points (Fig. 2B-i), consistent with near-steady-state cycling, where the fraction of cells observed in each phase reflects the relative time that cells spend in that phase across the population, which indicates ergodicity. Normality (p=0.822) and variance (p=0.418) tests were passed; two-way ANOVA with Tukey correction confirmed statistical differences only between phases within each condition (*p < 0.05). The average cell area trended downward from early to late timepoints but without significance: early G1 2039 ± 228 µm² vs. late G1 1802 ± 184 µm²; early S/G2/M 2257 ± 178 µm² vs. late S/G2/M 2025 ± 139 µm² (normality p=0.492, variance p=0.907; no factor-level effect, Fig. 2B-ii). Tubulin spread was consistently higher in S/G2/M than in G1 but declined between early and late: early G1 976 ± 82 µm² vs. late G1 655 ± 110 µm²; early S/G2/M 1216 ± 117 µm² vs. late S/G2/M 718 ± 62 µm² (Fig. 2B-iii). These data suggest that tubulin remodeling is the most sensitive morphological feature to the passage of time in culture.

**Figure 2.**
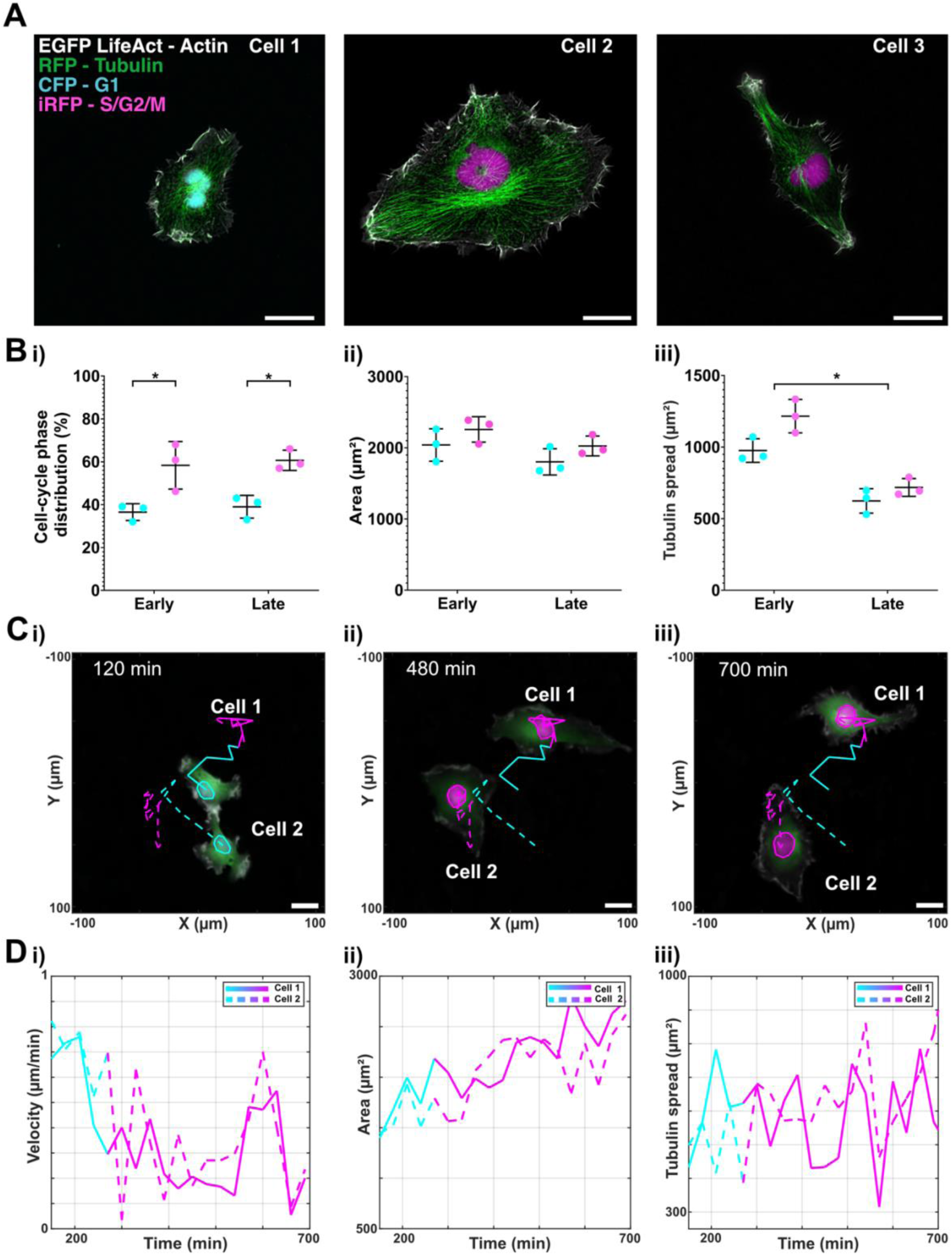
CC-aware phenotyping of HT1080 CALIPERS in free space. (A) Representative four-channel fluorescence images of HT1080 CALIPERS cells in G1 (CFP, Cell 1) and S/G2/M (iRFP, Cells 2–3) with actin (EGFP LifeAct) and tubulin (RFP). Scale bars, 25 µm. (B) Quantification of static phenotyping at early (4 h, n = 60 cells) and late (24 h, n = 82 cells) time points. (i) CC phase distribution: 37% ± 4% G1 vs. 58% ± 11% S/G2/M (early) and 39% ± 5% G1 vs. 61% ± 5% S/G2/M (late). (ii) Cell area: early G1 2039 ± 228 µm², S/G2/M 2257 ± 178 µm²; late G1 1802 ± 184 µm², S/G2/M 2025 ± 139 µm². (iii) Tubulin spread early G1 976 ± 82 µm², S/G2/M 1216 ± 117 µm²; late G1 655 ± 110 µm², S/G2/M 718 ± 62 µm². *p < 0.05 between phases within conditions (two-way ANOVA; time/area effect n.s.). (C) Dynamic phenotyping tracks of two cells across 700 min showing G1-S/G2/M transitions. (D) Dynamic quantification of (i) velocity (average 0.35 µm/min, mean path length 192 ± 22 µm, n = 53 tracks), (ii) area (increase toward S/G2/M), and (iii) tubulin spread (high variability, frequent 1.5-fold changes within 1 h).

**Figure 3.**
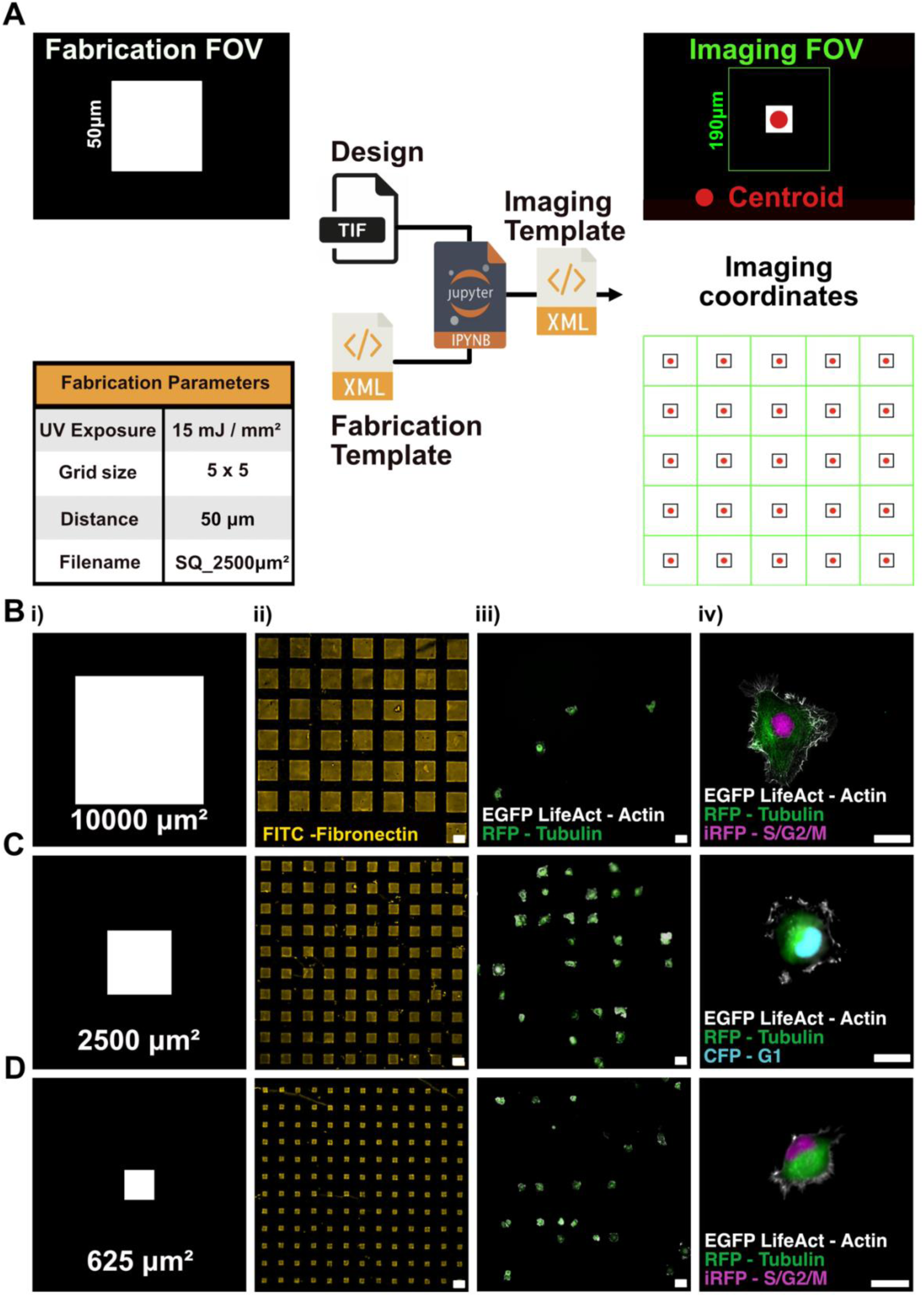
Fab2Mic pipeline for correlative fabrication-to-microscopy of engineered ECM islands. (A) Schematic of Fab2Mic workflow translating fabrication layouts (.tif + .xml) into imaging coordinates (.xml). Example shows a 50 µm square mask (SQ_2500 µm², 5 × 5 array) with centroids (red) and imaging FOV (green). (B–D) Representative binary masks (i), FITC-fibronectin arrays imaged at 477 nm (ii), HT1080 CALIPERS seeded on islands (iii), and high-resolution 100× imaging of single cells (iv) for engineered ECM islands of 10000 µm² (B), 2500 µm² (C), and 625 µm² (D). Arrays fabricated as 25×25 grids with 50 µm spacing. High-resolution imaging shows actin (EGFP LifeAct), tubulin (RFP), and nuclei labeled with CFP (G1) or iRFP (S/G2/M).

To capture how migration evolves through phases, we performed long-term CC-aware live imaging for 15 h at 30-min cadence (three independent experiments, ≥3 FOV each). Representative trajectories (Fig. 2C) show two cells tracked across CFP-iRFP transitions. Overall, across 54 free-space tracks, cells spent on average 4.5 ± 2.0 h in G1 and 5.2 ± 2.0 h in S/G2/M, respectively. Velocity traces revealed an average velocity of 0.37 ± 0.20 µm/min in G1 and 0.32 ± 0.20 µm/min in S/G2/M, with a mean path length of 192 ± 22 µm. Instantaneous velocity peaked during G1, dropped at the G1-S/G2/M transition, and rose again before mitosis (Fig. 2D-i), suggesting that migration speed is coupled to CC-state, potentially reflecting higher motility during growth, reduced dynamics during DNA replication, and reorganization of the cytoskeleton before division. Area measurements increased towards S/G2/M, consistent with static results (Fig. 2D-ii). Tubulin spread fluctuated substantially, often changing 1.5-fold within an hour, reflecting intracellular remodeling which could not be captured from static snapshots (Fig. 2D-iii). A small G0/G1-arrested subpopulation was present but excluded from this analysis, as arrested cells remain outside the proliferative cycle and thus do not engage in the dynamic remodeling observed in cycling populations; these are addressed in Fig. 4.

**Figure 4.**
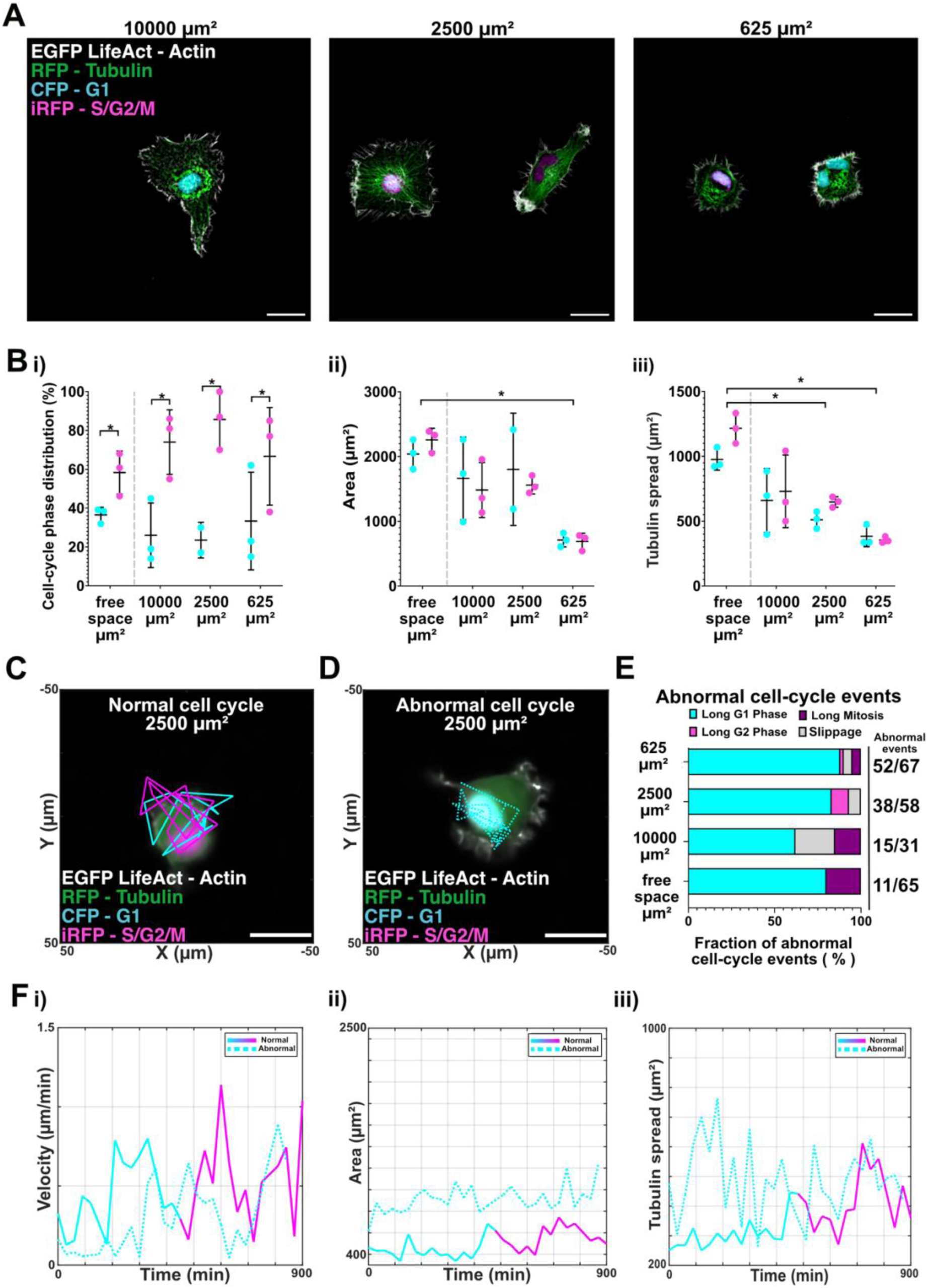
CC-aware phenotyping of HT1080 CALIPERS under confinement reveals continuous scaling of morphology and cell-cycle fidelity. (A) Representative images of HT1080 CALIPERS cells seeded on engineered ECM islands of 10000, 2500, and 625 µm², showing actin (EGFP LifeAct), tubulin (RFP), and nuclei in G1 (CFP) or S/G2/M (iRFP). Scale bars, 25 µm. (B) Static phenotyping across free space and confinement. (i) CC phase distribution: free space (40% G1, 60% S/G2/M) shifts under confinement, with G1 reduced to 26% ± 17% at 10000 µm² and 23.5% ± 9% at 2500 µm², with variability peaking at 625 µm² (33% ± 25%). (ii) Cell area decreased progressively with confinement, from >2000 µm² in free space to 711 ± 106 µm² at 625 µm². (iii) Tubulin spread followed the same trend, halving from 659 ± 246 µm² (G1, 10000 µm²) to 383 ± 80 µm² (G1, 625 µm²). *p < 0.05 between phases within conditions. (C–D) Representative dynamic trajectories at 2500 µm² show normal CC progression (C) and a prolonged G1 abnormality (Long G1) (D). (E) Frequency of abnormal CC events increased continuously with confinement: 17% in free space, 48% at 10000 µm², 66% at 2500 µm², and 78% at 625 µm². Long G1 phases dominated at small islands (75% of abnormal events at 625 µm²), while S/G2/M-G1 slippage emerged but decreased with island size (21% at 10000 µm², 5% at 2500 µm², and 4% at 625 µm²). Slippage was observed only under confined conditions and was interpreted as a confinement-associated vulnerability. Rare Long S/G2/M and Long Mitosis events were also observed, together accounting for about 5% of all abnormal events. Percentages were calculated over the total number of tracked cells pooled across three independent experiments. Cell death events were also recorded (5 in free space, 1 on 10000 µm² islands, 7 on 2500 µm² islands, and 9 on 625 µm² islands) but not included in the visualization for clarity of abnormal CC-phenotypes. (F) Dynamic comparisons at 2500 µm² between a normal and abnormal (Long G1) cell. (i) Instantaneous velocity was higher in normal G1 (0.44 ± 0.22 µm/min) than in Long G1 (0.29 ± 0.23 µm/min). (ii) Abnormal cells maintained 2× larger G1 area (912 ± 127 µm² vs. 465 ± 98 µm²). (iii) Tubulin spread expanded 2× during normal S/G2/M progression (309 ± 56 to 416 ± 98 µm²) but fluctuated broadly without net growth in Long G1 cells (489 ± 122 µm²).

Taken together, these results suggest that in free space, HT1080 CALIPERS cells maintained a near steady-state distribution of approximately 40% in G1 and 60% in S/G2/M, in line with previous reports^27,37,38^. Dynamic assays complement the picture by directly linking motility to cell-cycle transitions. Existing in vivo and ex vivo imaging studies^39^ demonstrate that CC–dependent regulation of invasion occurs. At the same time, in vitro longitudinal tracking approaches like ours^34,37,40^ are valuable as they can capture instantaneous velocity, trajectory, and intracellular variability, inaccessible to static endpoints.

### Implementation and characterization of a correlative pipeline for photofabrication and imaging

Having characterized our HT1080 CALIPERS reporter line in free space, we set out to study how cell migration and morphology evolve under confinement, by engineering adhesive ECM patterns via photofabrication that constrain cell migration over time. We realized that manually performing photofabrication and imaging as separate steps was both error-prone and inefficient. Mapping fabricated arrays and imaging fields-of-view (FOVs) via manual trial-and-error could lead to inaccuracy, which wasted imaging or analysis time. Instead, we developed Fab2Mic, a correlative fabrication-to-microscopy pipeline that directly translates photofabrication layouts into imaging coordinates, ensuring efficient and reproducible alignment between fabrication and live imaging. (Fig. 3A).

The algorithm takes the fabrication design (.tif) and metadata (.XML) as input, extracts feature centroids and array geometry and exports an XML file that can be used immediately for multi-point acquisition in most microscopes via commercial software or MicroManager. Smaller features, less than 500 µm in size, are centered in the imaging FOV, whereas larger features are tiled seamlessly to minimize redundancy. We validated this method on a 50 µm square design (5 × 5 array, 15 mJ/mm² exposure, 50 µm spacing) and confirmed that centroids (red) and imaging FOVs (green) matched the fabricated array on inspection (Fig. 3A). As a proof-of-concept, we also fabricated a 2D grayscale rendering of a picture of the Ponte Coperto in Pavia, coated it with FITC-fibronectin, and imaged a composite of 4×3 stitched FOVs (Fig. S2A-B, Movie S1). The Fab2Mic routine automatically generated the coordinate map and enabled acquisition without any manual adjustment.

We then applied this pipeline to fabricate engineered ECM islands of three sizes (10000, 2500, and 625 µm²) arranged as 25 × 25 arrays with a 50 µm spacing (Fig. 3B–D). Widefield 477 nm imaging of FITC-fibronectin confirmed sharp island boundaries and reproducible array geometry. Because single-cell occupancy is essential for unbiased phenotyping, we adopted a probabilistic seeding strategy that deliberately favors single-cell islands. On average, more than 50% of the occupied islands contain a single cell, although this comes at the cost of an overall occupancy of about 20% of the available islands (Fig. S2D–E). This trade-off was strategic: by combining probabilistic seeding with the automated Fab2Mic pipeline, we ensured reproducible access to single-cell islands while substantially reducing manual settings and registration time. After seeding HT1080 CALIPERS cells at optimized densities (4× array size for 625 and 2500 µm², 2× for 10000 µm²), cells adhered selectively to the islands. Finally, high-resolution imaging centered on individual islands revealed actin filaments aligned with fibronectin boundaries, microtubule networks spanning the cytoplasm, and CC phases marked by CFP (G1) or iRFP (S/G2/M).

Taken together, these results validate Fab2Mic as a correlative pipeline that reduces alignment errors in multipoint automated acquisitions, enables multiplexed imaging, and streamlines assay execution by saving time. By demonstrating robust single-cell occupancy across engineered ECM islands of decreasing size, we established the feasibility of combining photofabrication and CC-aware live imaging for quantitative studies of confined migration.

### Quantifying confinement effects on CC progression and proliferation

The combination of a validated reporter line, free-space baselines, and the Fab2Mic pipeline for high-content correlation between fabrication and imaging enables the systematic study of the influence of confinement on both migration and proliferation. Using engineered ECM islands of 10000, 2500, and 625 µm², we quantified CC distribution, morphology, and cytoskeletal dynamics under defined geometric constraints, comparing directly against free-space conditions. From the outset, imaging was restricted to islands occupied by single cells, ensuring unbiased phenotyping at the individual cell level.

Cells adapted to the adhesive island boundaries based on their free-space area: cells retained large spread morphologies on 10000 µm² islands (roughly five times their free space area), while at 625 µm² the cytoskeleton filled the available area (Fig. 4A). Interestingly, the CC distribution and cytoskeletal architecture show a continuous scaling in confinement (Fig. 4B). This scaling suggests that confinement acts as a continuum rather than a binary on/off state, with gradual modulation of both morphology and CC fidelity. In free space, approximately 40% of cells were in G1 and about 60% in S/G2/M. On the engineered ECM islands, the G1 fraction decreased to 26% ± 17% (10000 µm²) and 23.5% ± 9% (2500 µm²), with partial recovery to 33% ± 25% at 625 µm², suggesting a relative accumulation in pre-mitotic phases under confinement. Notably, variability increased with confinement, particularly in G1, where standard deviation rose to ±25% at 625 µm² (Fig. 4B). Cell area followed a similar trend: from over 2000 µm² in free space to 1663 ± 639 µm² (G1) and 1483 ± 425 µm² (S/G2/M) at 10000 µm², 1803 ± 867 µm² and 1559 ± 137 µm² at 2500 µm², and just 711 ± 106 µm² and 688 ± 128 µm² at 625 µm². Tubulin spread also decreased with confinement, roughly halving from 659 ± 246 µm² (G1, 10000 µm²) to 383 ± 80 µm² (G1, 625 µm²). Dynamic phenotyping revealed how these morphological effects intersect with CC fidelity (Fig. 4C–F, Fig. S5, Movie S2).

To test whether the reduced G1 fraction observed in static imaging reflected an artifact, we performed dynamic imaging as described in Fig. 2 and evaluated cells with normal and abnormal CC events separately. Across 156 tracked cells (31 at 10000 µm², 58 at 2500 µm², 67 at 625 µm²), 42 cells exhibited normal CC progression (Fig. S4), with average phase durations of 39% G1 and about 61% S/G2/M, even under confinement. However, abnormal CC events rose with increasing confinement, from a baseline of 17% in free space to 78% at 625 µm² (Fig. 4E). Abnormal events were classified as Long G1, Long S/G2/M, Long Mitosis, or S/G2/M-G1 slippage (see classification of abnormal cell-cycle in Methods section). While Long G1 and S/G2/M-G1 slippage dominated under confinement, rare instances of Long S/G2/M and Long Mitosis were also observed, together accounting for approximately 5% of all abnormal events. Long G1 phases dominated confinement-induced anomalies, increasing from 37% of abnormal events in free space to 75% at 625 µm². Conversely, S/G2/M-G1 mitotic slippage emerged^41,42^, but its incidence decreased with tighter constraints (21% at 10000 µm² to 4% at 625 µm²). In the context of this study, S/G2/M-G1 slippage was observed only under our confined conditions and was interpreted as a confinement-associated vulnerability. Representative trajectories at 2500 µm² (Fig. 4C–D), similar to the projected area of cells in free space, highlight the divergence: a normal cell progressed from G1 to S/G2/M with G1 velocity averaging 0.44 ± 0.22 µm/min, while a Long G1 cell had a lower motility (0.29 ± 0.23 µm/min). Morphologically, abnormal G1 cells maintained 2× larger areas (912 ± 127 µm² vs. 465 ± 98 µm²) and broader tubulin spread (489 ± 122 µm² vs. 309 ± 56 µm²), lacking the 2× cytoskeletal expansion observed in normal S/G2/M transitions (416 ± 98 µm²).

Taken together, these results indicate that confinement continuously modulates both morphology and CC fidelity, consistent with prior reports^7,19,43^ reduced adhesive area compresses cytoskeletal assembly. It increases the frequency of abnormal phenotypes, particularly prolonged G1 states (Long G1)^44^. Notably, while static imaging suggested a reduced representation of normal G1 under confinement, dynamic imaging revealed two distinct populations: normal and abnormal cycling cells. This indicates that ergodicity assumptions must be applied with caution. Automated cell tracking and cell-cycle phase assignment, as the one used here, provide critical value in distinguishing these subpopulations.

## Discussion

This study establishes a vertically integrated platform that unifies multiplexed CC reporters, engineered ECM islands, and correlative high-content imaging to resolve the coupling between migration and CC under geometric confinement. By combining spectral re-allocation through FUCCIplex (freeing the green/red channels for structural phenotyping), bridging the registration–throughput gap with Fab2Mic, and enabling longitudinal single-cell tracking, we convert confinement into a tunable parameter, consistent with our results. Indeed, as the adhesive area decreases, the G1 fraction declines, cytoskeletal organization becomes compressed, and variability increases. Embedding CC resolution directly into high-content morphological analysis provides a practical framework for systematic phenotyping of migration and proliferation under constraint, extending classic adhesion-^22,23^ and geometry-dependent control of proliferation^17^ including nuclear volume–G1 coupling on micropatterns^25,26^. Our results also support the importance of including live-cell imaging experiments to provide dynamic phenotyping, which revealed single-cell confinement-associated behaviors invisible to static imaging: including Long G1 or Long S/G2/M phases and S/G2/M–G1 mitotic slippage. These phenotypes are consistent with reports that mechanical stress impairs mitotic progression and promotes slippage-like outcomes under constraint^42,45,46^

More broadly, our platform extends the pioneering framework established by Ingber, who demonstrated how adhesion geometry and cytoskeletal tension regulate proliferation^17^. By integrating CC resolution through FUCCIplex and confinement control via Fab2Mic, we embed this principle into a high-content, imaging-based screening modality. This enables systematic interrogation of how physical context shapes proliferative fate decisions. The approach is compatible with large-scale image-based screens, where CC-aware analysis can be applied to evaluate compound effects on CC-migration coupling, or to design microenvironments that guide morphogenesis in engineered tissues^47^. Time-resolved high-content screening workflows have precedent (e.g., genome-scale live-cell imaging of mitosis), demonstrating the feasibility and value of temporal phenoprints^48–50^. However, scaling dynamic, multi-channel assays demands strengthened quality control. Scaling such assays to true high-throughput, however, requires strengthening the quality-control framework. Segmentation errors, tracking identity swaps, and channel registration drift remain partial bottlenecks, and manual curation was still necessary in this study for a subset of the data (∼20%). Future integration of AI-based segmentation and standardized quality controls (QC) benchmarks (e.g., error rates, reproducibility metrics) will be essential to translate CC-aware assays into robust pharma-ready pipelines, including multi-metric artifact detection and image-level QC (focus/illumination/saturation checks), which are being democratized via open guidance/toolkits for large-scale screens^51^. For example, cell-level QC workflows utilize clustering coupled with one-class support vector machines (SVMs) to flag segmentation artifacts and salvage valid cells from partially corrupted images^52^. Regardless of these future improvements, we believe our work bridges classical mechanobiology with next-generation tissue engineering, providing both a conceptual extension and a practical toolkit for probing how cells “read” geometry and confinement to balance migration and growth.

In the cancer context, our study contributes by introducing a 4-color reporter HT1080 line that allows systematic exploration of CC-migration coupling. We deliberately selected conditions that emphasize long-term behaviors, whereas actual metastatic migration in vivo is typically a transient, short-lived event. The value of our approach lies less in reproducing metastatic dissemination per se and more in providing a robust system to study how CC dynamics interact with migratory states. This perspective aligns with the “go-or-grow” framework^22,40,53^, and with pathology observations that invasive fronts are often slow-cycling while cores are proliferative^24,54,55^. In vivo, metastatic cells “go” to a distant site and then “grow,” whereas in our system, cells “go” but remain confined, leading to gradual accumulation of CC stress. Although caution is warranted when extrapolating to clinical applications, the CC abnormalities revealed under prolonged confinement demonstrate that cells become vulnerable to further proliferation, a state that, in principle, could prevent the development of a large metastatic mass. By contrast, recent findings show that confinement-induced states can also be more drug-tolerant, as HMGB2-high melanoma cells under mechanical confinement resist Taxol and BRAF/MEK inhibitors^54^. Reconciling these observations highlights a critical dependency on therapeutic context: most cytotoxic drugs preferentially target rapidly proliferating cells, so confinement-driven cell-cycle exit may simultaneously suppress metastatic outgrowth (beneficial) but blunt the efficacy of conventional chemotherapy (detrimental). Mechanistically, Taxol triggers mitotic arrest; therefore, non-dividing, confined cells are inherently less affected, and confinement-driven HMGB2 programs slow cycling and tolerance, with HMGB2 overexpression impairing BRAF/MEK response in vivo^7^. This tension highlights the importance of considering CC-migration coupling within the broader “go-or-grow” framework and evaluating confinement as both a source of mechanistic vulnerability and a driver of therapy resistance. Notably, some in vitro studies do not observe a strict trade-off: FUCCI-based analyses reported comparable motility in G1 vs S/G2/M and even under pharmacologic arrest^40^, and cross-line surveys found migration can correlate with proliferation in certain cancers^53^. Our results suggest that the appropriate geometric confinement can be leveraged to play along the go-or-grow continuum. This framing opens the door to thinking of designing confinement with a therapeutic goal. Analogous to immuno-engineered hydrogels designed to capture or prime immune cells^56–58^, one could envision biomaterials tailored as exploratory “capture sites” for metastatic cells. While speculative, this concept illustrates how our vertically integrated platform for CC-migration coupling assessment may inform both mechanistic mechanobiology studies as well as exploratory therapeutic screenings.

## Material and methods

### Cell line generation

HT1080 human fibrosarcoma cells (Ibidi, #HT-1080-LifeAct-TagGFP2, catalog# 40101) were genome-engineered to generate a four-color CALIPERS reporter line.

#### CRISPR/Cas9 Tubulin tagging

Endogenous α-tubulin (TUBA1B, NM_006082.3) was tagged with tagRFP using the Thermo Fisher Scientific TrueTag system. The donor sequence and primers with locus-specific homology arms for RFP tagging of the N-terminus of α-tubulin (TUBA1B, NM_006082.3) were designed with the assistance of the TrueDesign Genome Editor tool (Thermo Fisher Scientific)^59^. The linear TrueTag dsDNA donor was amplified by PCR using Phusion Flash High-Fidelity PCR Master Mix (TrueTag Donor DNA Kit, RFP, Thermo Fisher Scientific, catalog# A42993). The ribonucleoprotein (RNP) complex was assembled by combining TrueGuide synthetic sgRNA (Thermo Fisher Scientific; target sequence: GCACGGCTTACTCACCATAG) with TrueCut Cas9 Protein V2 (Thermo Fisher Scientific, catalog# A36496). TrueCut Cas9 Protein V2 (2 µg, 12 pmol), sgRNA (400 ng, 12 pmol), and the TrueTag donor (500 ng) were resuspended in 10 μL of Resuspension Buffer R (Thermo Fisher Scientific, catalog# MPK1025) and subsequently mixed with 8 × 10^4^ HT1080 cells resuspended in 8 μL of the same buffer, yielding a final reaction volume of 18 μL. A 10 μL aliquot of this mixture was electroporated using the Neon Transfection System (Thermo Fisher Scientific, catalog# NEON1S) with the following parameters: 1200 V, 20 ms, 4 pulses. Following electroporation, cells were seeded into 24-well plates containing 0.5 ml of Dulbecco’s Modified Eagle Medium / Ham’s F-12 Nutrient Mixture without phenol red (DMEM F-12, Gibco, catalog# 21041-025), supplemented with 10% heat-inactivated fetal bovine serum (FBS, Gibco, catalog# 10270-106) Cultures were incubated at 37 °C in a humidified CO_2_ atmosphere. RFP-positive cells were subsequently isolated and expanded in a clonal manner. Edited clones were validated via live-cell fluorescence microscopy and PCR (Fig. S1A, C)

#### FUCCIplex sensor integration

The FUCCIplex cassette was derived from pBOB-EF1-FastFUCCI-Puro (Addgene, plasmid #86849) by replacing mKO2 (Cdt1 degron) with mTurquoise2 and hmAzami Green (Geminin degron) with miRFP670. The cassette was cloned into a third-generation lentiviral backbone (pLV[Exp]-Hygro-EF1A, VectorBuilder, catalog# VB210825-1211nbd) with hygromycin resistance. Lentiviral particles were produced in HEK293T packaging cells (ATCC, catalog# CRL-1573), seeded at 2.5 × 10⁶ cells per 10 cm dish pre-coated with 0.2% gelatin (Sigma-Aldrich, catalog# G1393). Transfection was performed with Lipofectamine 2000 (Thermo Fisher Scientific, catalog# 11668019). Viral supernatants were collected at 48 h, clarified by centrifugation, and applied to HT1080 cells. More in detail, HT1080 cells were plated at a density of 6 × 10⁴ cells per well in 12-well plates containing complete DMEM F-12 medium. After 24 h, the culture medium was replaced with fresh complete medium supplemented with 8 µg/mL polybrene (Sigma-Aldrich, catalog# TR-1003-G). Concentrated FUCCIplex lentiviral particles (>10⁸ TU/mL, VectorBuilder; 10 µL resuspended in 500 µL of complete medium) were then added directly to the wells. Cells were incubated overnight at 37°C in a humidified 5% CO_2_ atmosphere. The following morning, the viral medium was aspirated and replaced with fresh complete medium. After 4 days, positively transduced cells were selected using Hygromycin B (Invitrogen, catalog# 10687010, 20 µL/mL from a 50 mg/mL stock in PBS). Resistant populations were clonally seeded and expanded. PCR and confocal fluorescence imaging verified correct integration and reporter expression

### PCR and RT-qPCR

Genomic DNA was extracted from HT1080 EGFP LifeAct and FUCCIplex clones using the DNeasy Blood and Tissue Kit (Qiagen, catalog# 69504) according to the manufacturer’s instructions. For genomic PCR, two different primer pairs were used to amplify the TUBA1B WT locus and the RFP-tagged gene. Specifically, primer pair 1-3 recognizes the WT locus, while primer pair 2-3 generates an amplicon only in the condition where the gene has been edited. PCR products were analyzed by agarose gel (1%) electrophoresis. PCR was carried out using Platinum PCR SuperMix, High Fidelity (Thermo Fisher Scientific, catalog# 12532016).

Total RNA was isolated using Trizol reagent (Thermo Fisher Scientific, catalog# 15596026), treated with DNase I (Thermo Fisher Scientific, catalog# EN0521), and reverse-transcribed using SuperScript™ IV Reverse Transcriptase (Thermo Fisher Scientific, catalog# 18090200). RT-qPCR was performed with primers specific for miRFP670, Hygromycin, and FUCCIplex. Relative expression levels were determined using the ΔΔCt method, with HPRT as the internal control.

### Cell culture

HT1080 CALIPERS cells were maintained in Dulbecco’s Modified Eagle Medium / Ham’s F-12 Nutrient Mixture without phenol red (DMEM F-12, Gibco, catalog# 21041-025), supplemented with 10% heat-inactivated fetal bovine serum (FBS, Gibco, catalog# 10270-106) and 1% Penicillin– Streptomycin (Himedia, catalog# A001-100ML). Cells were cultured at 37 °C in a humidified 5% CO_2_ incubator and routinely split at 70% confluency. For experiment preparation, cells were detached using Trypsin/EDTA 0.25% (Thermo Fisher Scientific, catalog# 25200056), resuspended in complete DMEM F-12, and properly counted according to the experimental needs.

### Cytoplasmic vs. nuclear CFP/iRFP evaluation

To evaluate CFP and iRFP signal expression and localization, HT1080 CALIPERS were screened with high-resolution widefield fluorescence microscopy with a Nikon CFI Plan Apo Lambda S 40× silicone oil objective (Silicone oil immersion, NA 1.25, WD 0.30 mm, catalog# MRD73400). See the microscopy acquisition section for detailed imaging settings.

Samples were prepared by seeding 5 × 10^4^ cells on 35 mm #1.5 glass-bottom dishes (Ibidi, catalog# 81158) pre-coated with 20 µg/mL human plasma fibronectin (Sigma-Aldrich, catalog# F0895) in standard supplemented DMEM F-12 without phenol red. After 24 hours, the medium was refreshed, and the cells were imaged.

The dataset was manually curated to identify cells expressing either CFP or iRFP signals. For each population, signal localization was assessed to distinguish between exclusive nuclear versus cytoplasmic expression. GraphPad Prism (version 10.4.2) was used to generate the graphics of this panel set. Results are reported in bar graphs with mean ±SEM.

### Scraping and clonal seeding

To enhance the yield of nuclear iRFP localization of HT1080 CALIPERS, a gentle scraping method was employed. A total of 5 × 10^4^ cells were seeded on two 35 mm #1.5 glass-bottom dishes (Ibidi, catalog# 81158) pre-coated with 20 µg/mL human plasma fibronectin (Sigma-Aldrich, catalog# F0895) in standard supplemented DMEM F-12 without phenol red medium.

After 24 hours, the medium (DMEM F-12, Gibco, catalog# 21041-025) was refreshed before the imaging session. Widefield fluorescence microscopy was used to manually annotate regions of interest (ROIs) corresponding to nuclear iRFP-positive cells directly on the dish.

Subsequently, adherent populations were gently detached using a sterile cell scraper.

The scraped cell sample was resuspended in fresh medium and replated for expansion and subsequent clonal seeding. Once the population reached 70% confluency, cells were trypsinized with Trypsin/EDTA, resuspended in complete DMEM F-12 medium, counted, and diluted to a final concentration of ∼0.8 cells per well. Clonal seeding was performed in a sterile 96-well plate (Ibidi, catalog# 89626), where 100 µL was dispensed per well to favor single-cell deposition. To ensure optimal fluorescence imaging, 48 wells were seeded, avoiding the outer rows/columns of the plate. The 96-well plate was incubated under standard culture conditions (37°C, 5% CO_2_), and wells were monitored to confirm clonal origin.

Of the 48 initially seeded wells, 8 remained empty, leaving 40 available clones for evaluation. The medium was refreshed every other day with standard supplemented DMEM F-12 until day 7 of clonal expansion.

### Automated FUCCIplex clonal screening

A high-throughput live-cell imaging JOB was developed using Nikon NIS-Elements AR software (version 5.42.03)^60^ to scan a multi-well plate through a multichannel serial acquisition and large-image composition per well. The JOB wizard enabled selection of wells of interest, definition of imaging channel stacks, and specification of the reference channel for automated focus adjustment, which was based on contrast optimization.

The implementation utilized 638 nm for iRFP (S/G2/M), 546 nm for RFP (Tubulin), 477 nm for EGFP LifeAct (Actin, used as the reference plane), 446 nm for CFP (G1), and brightfield acquisition. Each well is acquired as a large image of a 6 x 6 field within a Nikon CFI Plan Fluor DIC 10× (air objective, NA 0.3, WD 16 mm, catalog# MRH00105). The entire well is then reconstructed by stitching with a 10% overlap and stored as a merged file.

### FUCCIplex selector routine

An automated FUCCIplex selector routine was developed using the General Analysis 3 module in Nikon NIS-Elements AR software (version 5.42.03)^60^. The routine was based on tubulin and FUCCIplex nuclear signals. Specifically, FUCCIplex nuclear signals were combined into a DAPI-equivalent (DAPIEq) mask that captured both CFP and iRFP expression. The mask generated from CFP and iRFP signals was created within thresholds defined by object intensity and size to exclude cytoplasmic signals. CFP-positive nuclei were selected with a mean intensity > 800 and nuclear diameter < 15 μm. iRFP nuclear-localized expression was selected with intensity > 600 and a nuclear diameter < 15 μm. Using the tubulin and DAPIEq masks, the algorithm considered the number of cells detected by tubulin as the ground truth. As a selection criterion, the DAPIEq object count was required to reach > 80% of the tubulin-based cell count for a well to be classified as an HT1080 CALIPERS four-reporter clone. The threshold was relaxed to 80% because early G1 or M-phase cells may be missed by the FUCCIplex selector but still appear in tubulin-based counts, leading to slight mismatches.

### Photofabrication

#### Substrate cleaning and passivation

35 mm #1.5 glass-bottom dishes (Ibidi, catalog# 81158) were washed in 2% Hellmanex III solution (Hellma Analytics, catalog# 9-307-0110), rinsed with DI water, and dried. Dishes were exposed to UV-ozone for 30 minutes (BioForce Nanosciences ProCleaner). Substrates were coated with 200 µL of 1 mg/mL Poly-L-Lysine (PLL, Sigma-Aldrich, catalog# P4707) for 30 minutes, rinsed with HEPES buffer (pH 8.3-8.6; Sigma-Aldrich, catalog# H3375), and incubated with freshly prepared 70 mg/mL PEG-SVA (Laysan Bio, catalog# SVA-PEG-5000) for one hour, followed by three DI water washes.

For #1.5 Ibidi 8-well glass plates (Ibidi, catalog# 80807), volumes were downscaled according to the single-well volume, and UV-ozone exposure was replaced with 200 μL of 1M NaOH treatment for 30 minutes, to restrict substrate functionalization to individual wells.

#### ECM islands and validation designs

Square engineered ECM island designs of 625 µm², 2500 µm², and 10000 µm² were generated in Inkscape (version 1.3)^61^ and converted into 8-bit grayscale format with a white square island foreground. Validation images to test the Fab2Mic routine were also generated to test both the small design with array repetition and a large photopatterned region. For validation, a large region was generated from an 8-bit grayscale.tif image of the “Ponte Coperto in Pavia”(Fig. S2A-B).

#### Photopatterning

35 mm round substrates were coated with a photoinitiator solution consisting of 66 µL ethanol (VWR, catalog# 20821.330) mixed with 4 µL PLPP gel (Alvéole, catalog# LPPG-100) and dried in the dark. For #1.5 8-well glass Ibidi plates, a surfactant solution (Surfactant Mix, Alvéole) was required to allow the photoinitiator to spread evenly across the entire well bottom and prevent meniscus formation. The solution was therefore prepared by mixing 97 % (v/v) DI water, 2.5 % PLPP gel, and 0.5 % Surfactant Mix. A volume of 150 μL was dispensed into each well and allowed to dry completely.

Photopatterning was performed using the PRIMO system (Alvéole)^29^ coupled to a Nikon Ti2 inverted epifluorescence microscope. The integrated 375 nm UV source and a Nikon CFI S Plan Fluor ELWD 20XC (air objective, NA 0.45, WD 6.9 – 8.2, catalog# MRH08230) provided illumination. Selected designs were projected through the optical path using a Digital Micromirror Device (DMD), enabling the transfer of user-defined designs onto the substrate with a resolution of 0.28 μm/px. Fabrication layouts were tuned by selecting the array size (5 × 5 or 25 × 25) and inter-pattern spacing (50 μm or 150 μm). Pattern alignment and exposure control were managed through the Leonardo software package (Alvéole, v.5.2)^62^.

#### ECM coating

Photopatterned substrates were incubated with 20 µg/mL FITC-fibronectin (Sigma-Aldrich, catalog# FNR01-A) in PBS for 30 minutes for validation tests, or with human plasma fibronectin (Sigma-Aldrich, catalog# F0895) for 10 minutes. Three PBS washes removed excess coating.

#### Cell seeding on ECM photopatterned islands

HT1080 CALIPERS cells were seeded on pre-coated engineered ECM islands with human plasma fibronectin (Sigma-Aldrich, catalog# F0895) at a density adjusted for single-cell occupancy: 4 × array size for 625/2500 µm² and 2 × array size for 10000 µm² patterns (Fig. S2C-E). Cells were dispensed in complete DMEM F-12 filled photopatterned Ibidi 8-well glass plates (Ibidi, catalog# 80807) or 35 mm #1.5 glass-bottom dishes (Ibidi, catalog# 81158). After 15 minutes of settling under the hood, cells were incubated at 37 °C and 5% CO_2_ for two hours before replacement with fresh phenol-red-free DMEM F-12 for imaging.

### Fab2Mic – Correlative fabrication-to-microscopy pipeline

A Python pipeline (Fab2Mic) was developed to translate photofabrication templates directly into imaging coordinates. The routine accepts as input a binary fabrication mask (.tif) and a photofabrication report (.XML). The XML file encodes essential metadata, including the scale factor, substrate position, grid size, spacing, and fabrication objective specifications. These parameters were parsed to reconstruct the layout and to generate coordinate maps compatible with microscope control software.

For high-resolution acquisitions (≤ 500 µm field of view), imaging positions were assigned to the centroid of each fabricated feature to preserve positional fidelity. For larger fields of view (> 500 µm), redundancy was minimized by computing a tiling grid, with fields automatically centered and spaced by their dimensions to ensure complete coverage without overlap (Fig. S2A-B). The pipeline, therefore, enables direct one-to-one registration between fabrication and imaging, scaling seamlessly from single-feature to array-level designs without manual intervention.

### Microscopy acquisitions

Acquisitions were performed on a widefield Nikon Ti2 inverted microscope with a CrestOptics X-Light V3 spinning disk confocal unit and an environmental chamber (Okolab Bold Line; 37 °C, 5% CO_2_, humidity). The microscope is equipped with a Nikon linear-encoder motorized stage with a Mad City Labs 100 µm range Z-piezo insert (Mad City Labs, catalog# NI-2-C312). Illumination was provided by the Lumencor Celesta light engine (405, 446, 477, 520, 546, 638, 740 nm; up to 800 mW), routed through multiband dichroics and appropriate emission filters.

The dichroic mirror wheel was selectively positioned with either a hard-coated Full Multi Band Penta CELESTA –DA/FI/TR/Cy5/Cy7-A (Nikon, catalog# MXR00543) or a Full Multiband dual CELESTA –CFP/YFP-A (Nikon, catalog# MXR00544). Depending on the excitation line, the beam was subsequently filtered through either a Multiband Full Penta FF01-391/477/549/639/741 (Semrock, catalog# FL-416877) or a Full Multiband Dual FF01-449/520 (Semrock, catalog# FL-411981).

Emission was collected using single-band filters mounted in the wheel: FF01-484/561 (Semrock, catalog# FL-412124) for CFP, FF01-685/40-25 nm (Semrock, catalog# FL-011482) for the iRFP signal, FF01-595/31 (Semrock, catalog# FL-004391) for RFP, or FF01-511/20-25 (Semrock, catalog# FL-004306) for EGFP.

Fluorescence was captured with the same Teledyne Photometrics Kinetix CMOS camera (6.5 µm pixels; 16-bit; native resolution 3200 × 3200 pixels, cropped to 2700 × 2700 pixels). Acquisition and hardware synchronization were controlled through NIS-Elements AR (v.5.42.03)^60^. Data were stored as .nd2 files.

#### Low-resolution clone selection

Static widefield and brightfield acquisitions were performed using a CFI Plan Fluor DIC 10× (air objective, NA 0.3, WD 16 mm, catalog# MRH00105). Clone selection was run with the custom automated CALIPERS clonal screening JOB in which each well was acquired as a large image of a 6 x 6 field within a CFI Plan Fluor DIC 10× (air objective, NA 0.3, WD 1.6 mm, catalog# MRH00105). Channel acquisition was sequentially ordered from 638 nm to 446 nm, with brightfield acquired last. The high-throughput JOBs covered 40 wells, with the shutter closed during stage movements. Illumination settings were:

● **iRFP (S/G2/M)**: 638 nm, exposure 300 ms, laser power 28.71 ± 3 mW
● **RFP (Tubulin)**: 546 nm, exposure 100 ms, laser power 15.79 ± 1.6 mW
● **EGFP (Actin)**: 477 nm, exposure 100 ms, laser power 15.21 ± 1.5 mW
● **CFP (G1)**: 446 nm, exposure 300 ms, laser power 3.12 ± 0.3 mW
● **Brightfield:** Dia lamp exposure 5 ms

#### Fab2Mic validation imaging

Fab2Mic validation was performed on square ECM designs of 625 µm², 2500 µm², and 10000 µm², as well as on a photopatterned image of the “Ponte Coperto in Pavia”. Designs were photopatterned and FITC-fibronectin-coated. Widefield fluorescence microscopy was used to validate the algorithm using NIS-Elements software^60^. Imaging coordinates were imported from the XML file returned by Fab2Mic. Acquisitions of the photopatterned FITC-fibronectin square arrays were performed with a Nikon CFI Plan Fluor DIC 10× (air objective, NA 0.3, WD 16 mm, catalog# MRH00105) and the 477 nm line set at laser power 60.2 ± 6 mW with a 30 ms exposure. Acquisitions of the “Ponte Coperto in Pavia” pattern were performed with a Nikon CFI Plan Apo 20× (air objective, NA 0.75, WD 1 mm, catalog# MRD00205) and the 477 nm line set at laser power 27.8 ± 2.8 mW with a 300 ms exposure.

#### High-resolution static confocal Z-stack

High-resolution static confocal imaging was performed with a Nikon CFI SR HP Plan Apo Lambda S 100× silicone oil objective (Silicone oil immersion, NA 1.35, WD 0.31 mm, catalog# MRD73950). Illumination settings were:

● **iRFP (S/G2/M)**: 638 nm, 2×2 binning, exposure 2 s, laser power 5.61 ± 0.6 mW
● **RFP (Tubulin)**: 546 nm, exposure 300 ms, laser power 1.8 ± 0.2 mW
● **EGFP (Actin)**: 477 nm, exposure 300 ms, laser power 1.9 ± 0.2 mW
● **CFP (G1)**: 446 nm, 2×2 binning, exposure 30 ms, laser power 1.4 ± 0.1 mW

Z-stacks were acquired using an MCL Piezo Z-device by capturing each channel across a 10 µm range within a step size of 0.3 μm

#### High-resolution static widefield imaging

Static widefield imaging was performed with a Nikon CFI SR HP Plan Apo Lambda S 100× silicone oil objective (Silicone oil immersion, NA 1.35, WD 0.31 mm, catalog# MRD73950), Nikon CFI Plan Apo Lambda S 40× silicone oil objective (Silicone oil immersion, NA 1.25, WD 0.30 mm, catalog# MRD73400) For 100× imaging, sequential four-channel acquisitions were carried out with the following settings:

● **iRFP (S/G2/M)**: 638 nm, exposure 50 ms, laser power 14.8 ± 1.5 mW
● **RFP (Tubulin)**: 546 nm, exposure 30 ms, laser power 6.7 ± 0.7 mW
● **EGFP (Actin)**: 477 nm, exposure 30 ms, laser power 5.9 ± 0.6 mW
● **CFP (G1)**: 446 nm, exposure 2 ms, laser power 0.8 ± 0.1 mW For 40× imaging, acquisitions were performed with:
● **iRFP (S/G2/M)**: 638 nm, exposure 100 ms, laser power 28.8 ± 3 mW
● **RFP (Tubulin)**: 546 nm, exposure 40 ms, laser power 14.8 ± 1.5 mW
● **EGFP (Actin)**: 477 nm, exposure 40 ms, laser power 13.5 ± 1.4 mW
● **CFP (G1)**: 446 nm, exposure 60 ms, laser power 2.2 ± 0.2 mW

#### Low-resolution dynamic live imaging

Long-term widefield acquisitions were performed using a Nikon CFI Plan Apo 20× (air objective, NA 0.75, WD 1 mm, catalog# MRD00205). Multipoint time-lapse imaging was conducted sequentially from 638 nm to 446 nm, with the shutter closed during stage movements.

For 20x imaging, sequential four-channel acquisitions were carried out with excitation settings as follows:

● **iRFP (S/G2/M)**: 638 nm, 4×4 binning, exposure 300 ms, laser power 21.4 ± 2.1 mW
● **RFP (Tubulin)**: 546 nm, exposure 50 ms, laser power 20.0 ± 2 mW
● **EGFP (Actin)**: 477 nm, exposure 50 ms, laser power 15.8 ± 1.6 mW
● **CFP (G1)**: 446 nm, 4×4 binning, exposure 20 ms, laser power 3.1 ± 0.3 mW

Time series spanned 15 hours with a frame interval of 30 minutes. Stage movements followed an optimized multipoint path. Nikon Perfect Focus remained engaged throughout overnight acquisitions to minimize drift. Samples were maintained in phenol-red-free DMEM F-12 medium.

#### Image preprocessing and visualization

Raw acquisitions were denoised and deconvolved in NIS-Elements using the Richardson–Lucy algorithm (10 iterations). A custom Fiji^63^ macro was applied for visualization, including a rolling-ball background subtraction (50 px radius) and channel-specific intensity level subtraction (446 nm = 50; 477 nm = 20; 546 nm = 20; 638 nm = 100). Scale bars were set at 25 µm.

### HT1080 CALIPERS CC-aware bioimage analysis

Static and dynamic microscopy datasets underwent a multi-step analysis pipeline to extract CC-aware dynamics, motility, and macro– and micro-scale morphology features.

#### Pre-processing

Low-resolution fluorescence acquisitions were first pre-processed by applying BaSiC flat-field correction^64^ to compensate for non-uniform illumination and background shading, yielding homogeneous image intensity. Denoising was performed using a Butterworth filter implemented in the *dexp* library (Royer Lab, GitHub)^65,66^. For all channels, parameters were calibrated on the first frame using calibrate_denoise_butterworth, and the resulting function was applied across all time points. Channel indexing and metadata were provided via a configuration file. Processed stacks were saved as 16-bit OME-TIFFs using *AICSImageIO*^67^. Image manipulations were performed with *scikit-image*^68^.

#### Segmentation and tracking

Nuclear masks were generated with StarDist (version 0.8.3, pre-trained fluorescent nuclei model)^31^. Channel signals were merged into a single DAPI-equivalent image, from which segmentation masks were obtained. The resulting DAPIEq masks were used as labels for motility analysis. Cytoskeleton masks were generated with Cellpose v2.0^32^ applied to RFP (tubulin) and EGFP LifeAct (Actin) channels. The segmentation derived from cytoskeletal signals defined macro-scale morphological labels for HT1080 CALIPERS. Tubulin microstructures were labeled with Fiji Otsu-thresholding^69^ on background-subtracted (rolling ball 50 px) and Gaussian smoothed (σ = 2 px) RFP (tubulin) channel (Fig. S3A).

#### FUCCIphase with morphology cues

Cell-cycle phase assignment was performed using FUCCIphase^34^, with newly calibrated intensity curves of the HT1080 CALIPERS cells. For each tracked nucleus, normalized CFP and iRFP mean intensities were measured within the DAPIEq mask. Phase classification was performed using fixed intensity thresholds derived from 24-hour reference recordings, yielding G1 (CFP-high/iRFP-low) and S/G2/M (iRFP-high/CFP-low). Intermediate G1/S events were grouped with S/G2/M for the analyses. Time-resolved motility and morphology were computed by linking nuclear centroids in Fiji^63^ using the TrackMate plugin^33^ with LAP tracking.

The nuclear signal intensity of the two FUCCI channels was used for static and dynamic CC-phase assignment, and the centroid of the nuclear labels was used for cell motility parameter computation. Additionally, CC-related nuclear morphology features were directly computed from the nuclear labels (Fig. S3B, Fig. S6). In parallel, static or time-resolved cytoskeletal features were extracted from cytoskeleton mask centroids within the same pipeline. Additional morphological features, such as perimeter, circularity, and ellipse aspect ratio (Ellipse AR), were evaluated (Fig. S3C, Fig. S7). The spread of tubulin microstructure was quantified through an automated pixel count macro applied to tubulin masks, with measurements converted to physical area (µm²).

For dynamic analyses, single-cell tubulin spread was computed per frame and paired with the concurrent FUCCIphase derived cell track. To link nuclear information with cytoskeletal information, a custom MATLAB script (FUCCIphase_morphology, version R2023b, 23.2)^70^ was developed to map each nuclear centroid to its enclosing cytoskeletal mask centroid within a 20 μm search radius, minimizing incorrect associations. The output was a unified FUCCIphase_morphology table (per-cell, per-frame for dynamic analyses) containing time stamps, positions, phase labels, cell-scale morphology, and tubulin spread. The same pipeline was applied to both static and dynamic datasets, acquired under both free-space and confined conditions.

### Static HT1080 CALIPERS CC-aware phenotyping

For free space condition, 5 × 10^4^ cells were seeded on two 35 mm #1.5 glass-bottom dishes (Ibidi #81158) pre-coated with 20 µg/mL human plasma fibronectin (Sigma-Aldrich, catalog# F0895) in standard supplemented DMEM F-12 without phenol red medium for Early (4-hour post seeding) vs Late (24-hours post seeding) timepoints evaluations. For confinement conditions, cells were seeded according to the engineered ECM island size and array dimension. See the photofabrication section for details on engineered ECM island seeding. In both conditions, the medium was refreshed two hours before the imaging session. Three independent high-resolution widefield fluorescence experiments were performed with a CFI SR HP Plan Apo Lambda S 100× silicone oil objective (Silicone oil immersion, NA 1.35, WD 0.31 mm, catalog# MRD73950).Static datasets were analyzed with the HT1080 CALIPERS CC-aware bioimage analysis pipeline and stratified by cell-cycle phase (G1 = CFP; S/G2/M = iRFP) to extract principally: average cell-cycle phase distribution, cell area, and tubulin spread per experiment (Fig. 2B, Fig. 4B). Additional information on nuclear morphology and cytoskeleton assembly is also quantified (Fig. S3).

### Dynamic HT1080 CALIPERS CC-aware phenotyping

For the free space condition, 5 × 10^4^ cells were seeded on 35 mm #1.5 glass-bottom dishes (Ibidi #81158) pre-coated with 20 µg/mL human plasma fibronectin (Sigma-Aldrich, catalog# F0895) in standard supplemented DMEM F-12 without phenol red. For confinement conditions, cells were seeded according to the square ECM size and array dimension. See the photofabrication section for details on ECM island seeding. In both conditions, the medium was refreshed two hours before the imaging session in each case. Three independent dynamic low-resolution fluorescence experiments were performed with a CFI Plan Apo 20× (air objective, NA 0.75, WD 1 mm, catalog# MRD00205). Dynamic datasets were analyzed with the HT1080 CALIPERS CC-aware bioimage analysis pipeline. CC-labeled tracks were computed using a custom MATLAB script (v.R2023b, 23.2)^70^ that generated color-coded CC on fluorescence images and extracted CC-aware motility metrics over time, including velocity, cell area, and tubulin spread (Fig. 2D, Fig. 4D). Comprehensive CC-related nuclear and cytoskeleton morphology features are evaluated (Fig. S6, Fig. S7). Additionally, cell-cycle phase duration, total path length, and net displacement stratified by cell-cycle phase (G1 = CFP; S/G2/M = iRFP) were quantified.

### Abnormal cell-cycle events analysis

Cell-cycle abnormalities were manually annotated across the three analyzed experiments of dynamic HT1080 CALIPERS CC-aware phenotyping. Five categories were defined. To classify abnormal CC events, ‘Long G1’ (>9 h) and ‘Long S/G2/M’ (> 10h) were pragmatically defined using a distribution-based threshold, set at two standard deviations above the mean of the free-space distribution. This approach follows precedent from single-cell timing studies that applied distribution thresholds for outlier detection^37^. Additional abnormal events included ‘Long Mitosis’, defined as persistence in M phase beyond the expected one-hour duration. Finally, rare events of direct S/G2/M-G1 re-entry without mitosis (‘slippage’) were identified exclusively under confinement. To minimize the risk of annotation errors, trajectories were manually curated, and only cases confirmed by visual inspection were retained.

### Statistical analysis

All statistical analyses were performed using GraphPad Prism v10.4.2^71^ and SigmaPlot v14^72^. Normality was assessed with the Shapiro–Wilk test, and equal variance with the Brown-Forsythe test. For parametric data, two-way ANOVA with Tukey’s post hoc correction was applied; for non-parametric distributions, Kruskal–Wallis tests were used. All static and dynamic phenotyping assays were performed in three independent experiments (n = 3), each derived from separate cell passages and sample/substrate preparations, representing both biological and technical replicates. Unless noted, values are reported as mean ± SD. Exceptionally, for cytoplasmic versus nuclear CFP/iRFP localization, values were averaged per field of view (FOV) and reported as mean ± SEM across FOVs. For abnormal CC event frequencies (Fig. 4E), percentages were calculated over the total number of tracked cells pooled within each confinement condition. A threshold of p < 0.05 was considered statistically significant. Graphical annotations in figures indicate significant comparisons (*p < 0.05).

### Author contributions

M.P. and F.S.P. conceived and demonstrated the concept of a vertically integrated system for cell-cycle-aware phenotyping under engineered confinements. M.P. and M.D. conducted the experiments and managed cell handling. M.D. performed the HT1080 genome editing to generate the CALIPERS line and carried out PCR experiments and analyses. M.P. performed fluorescence microscopy screening of CFP/iRFP expression and localization and subsequent studies. M.P. and M.D. carried out cell scraping and clonal seeding. M.P. developed the CALIPERS rescue pipeline, implemented the automated CALIPERS clonal screening JOB and FUCCIplex selector routine, and established clone E4 as the working line. M.P. designed the engineered ECM islands, performed substrate functionalization and photopatterning, and developed the correlative fabrication-to-microscopy (Fab2Mic) pipeline. M.P. executed all imaging experiments and analyzed the data using FUCCIphase (J.Z.’s software published in CALIPERS^34^) and the FUCCIphase_morphology script with additional morphological cues. E.T., J.Z., S.R., and A.E. supervised experiments and contributed to manuscript preparation. M.P. and F.S.P. co-wrote the initial manuscript. All authors discussed the results and provided feedback on the final version.

## Acknowledgements

This work was supported by the European Research Council (ERC) Starting Grant No. 852560 to F.S.P., by the Marie Sklodowska-Curie Postdoctoral Fellowships (HORIZON-MSCA-2023-PF, No. 101153603) awarded to A.E., and by the Italian Ministry of Education, University, and Research (MIUR) (FARE2020, Grant No. R20ZE54CTK) to F.S.P. ChatGPT (OpenAI, GPT-5, 2025) was used to refine phrasing, improve readability, and code formatting; all scientific content and analyses were developed and validated by the authors.

## Data and code availability

Representative raw microscopy datasets, processed outputs, generated tables, and custom analysis code supporting the findings of this study are available in <u>Zenodo</u>. Standard software packages used for image processing and quantification are described in the Methods section. Additional materials are available from the corresponding author upon request.

## Ethics statements

This study did not involve human participants, human data, or animal experiments. All experiments involving established cell lines were performed in accordance with institutional biosafety guidelines.

---

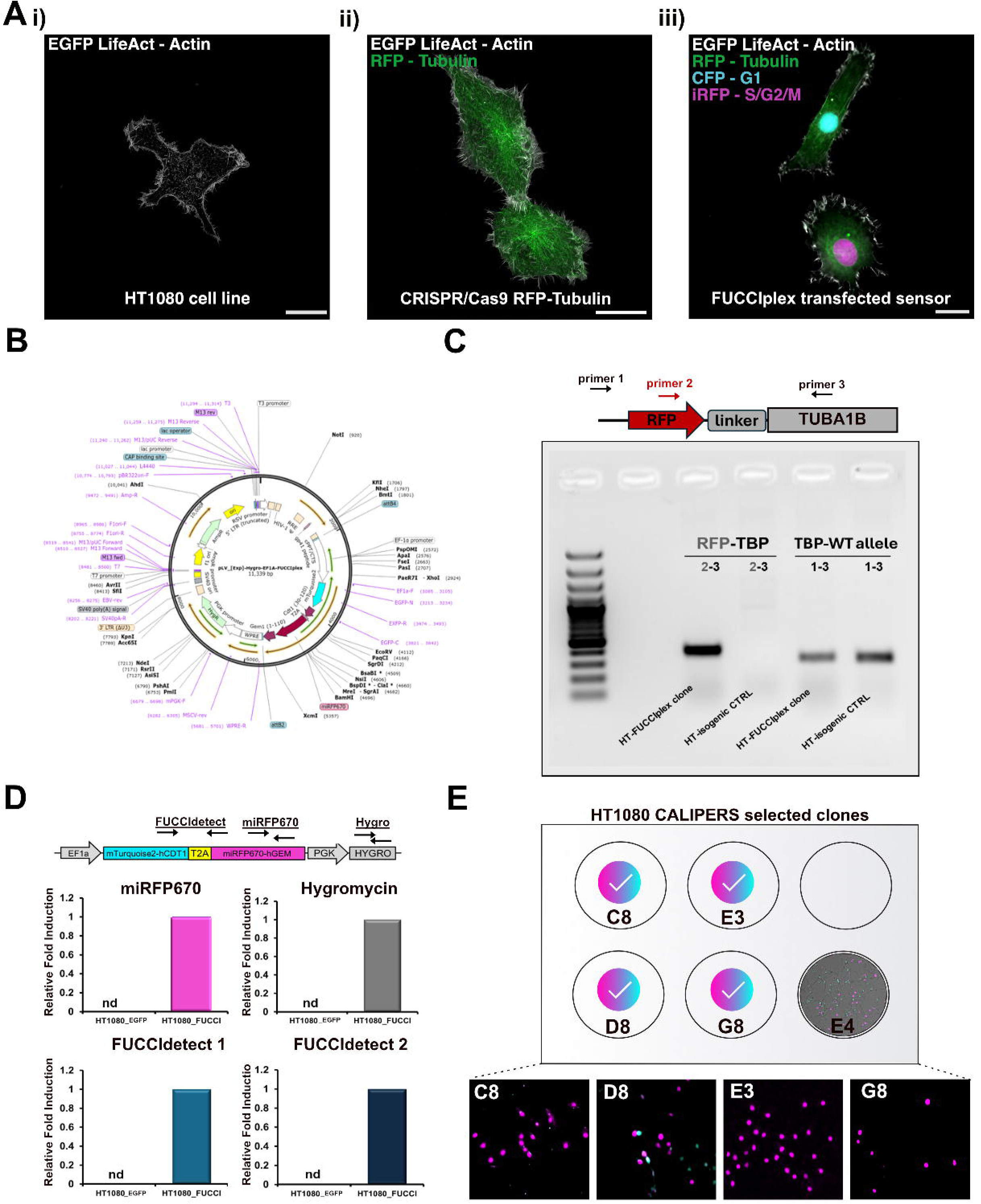

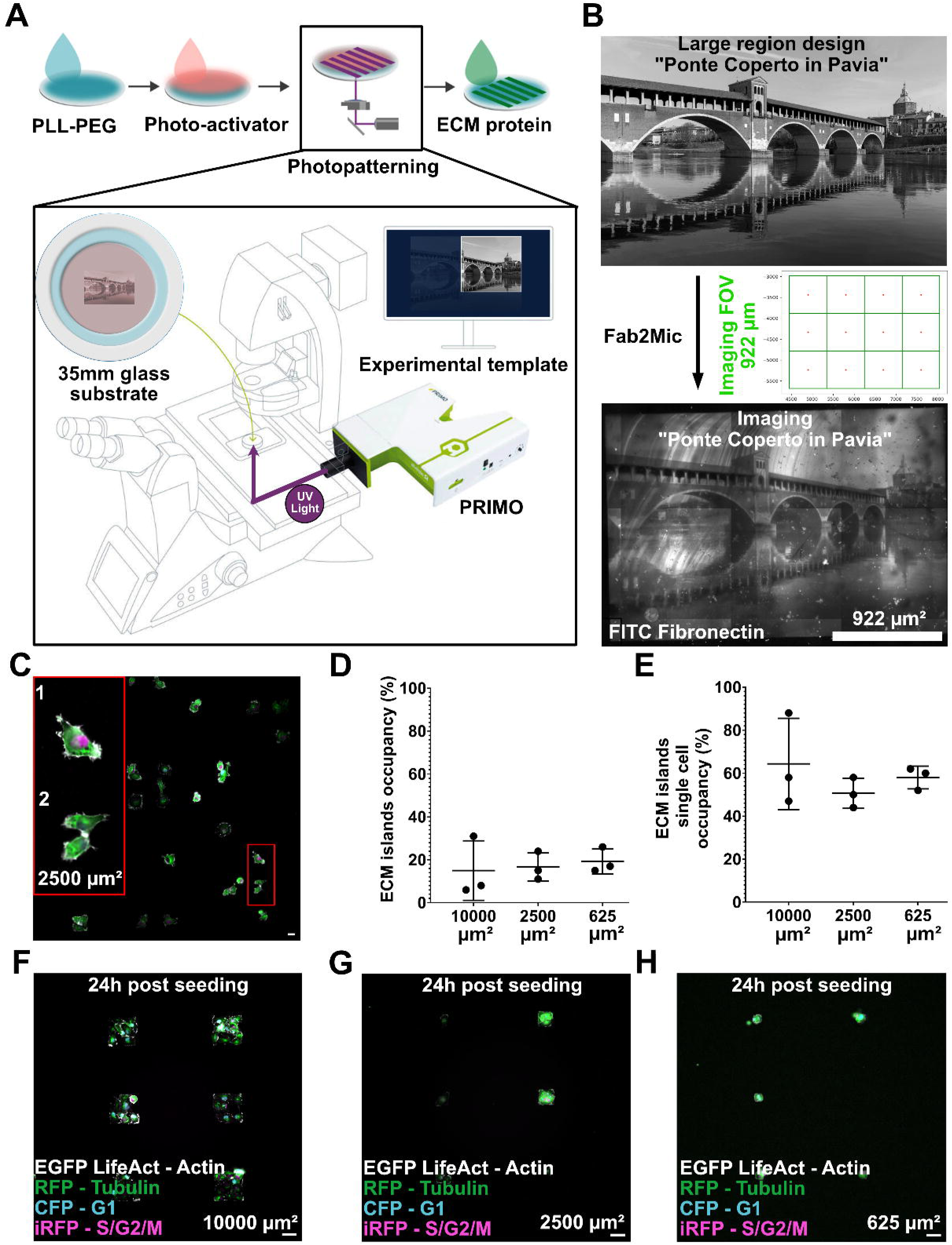

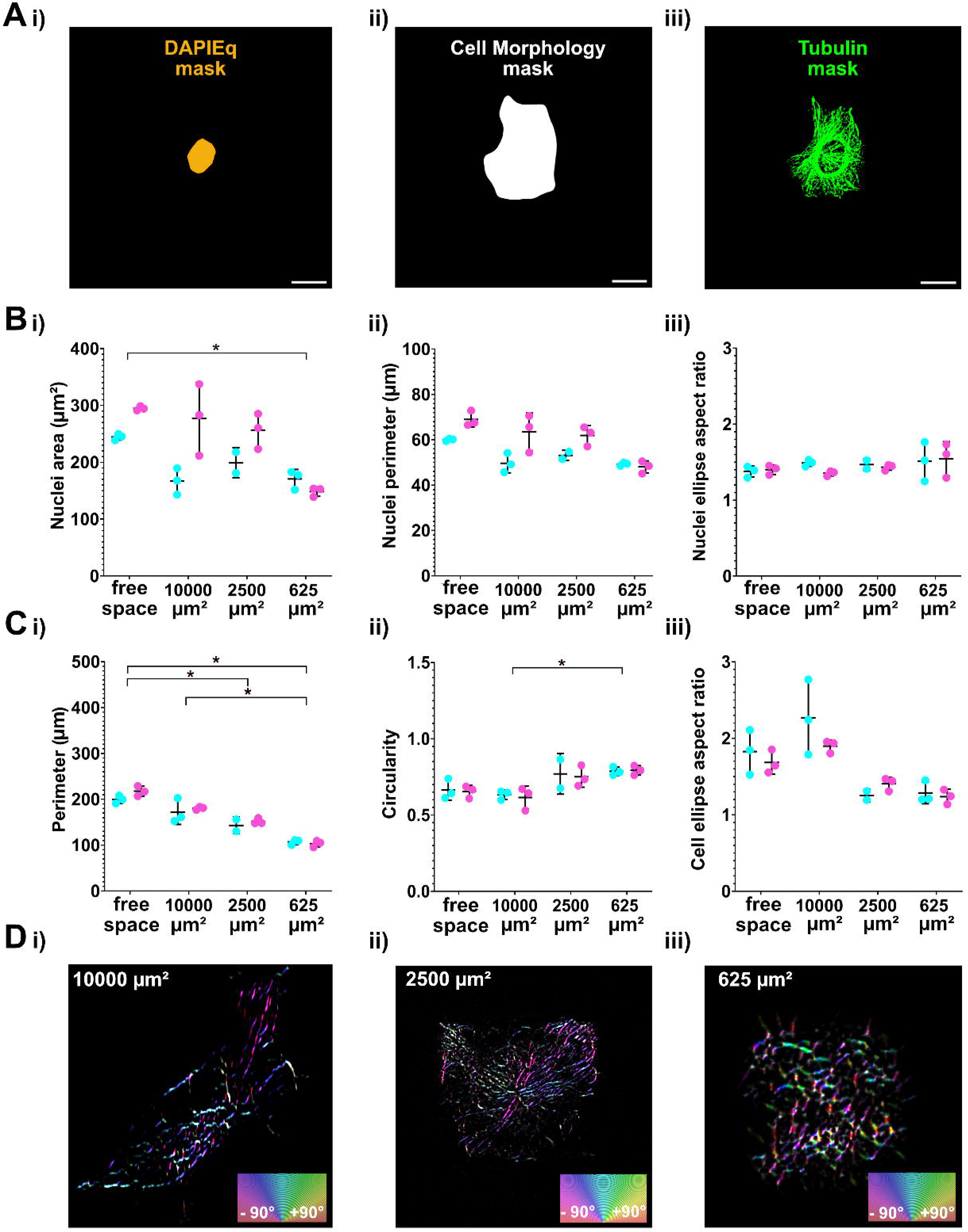

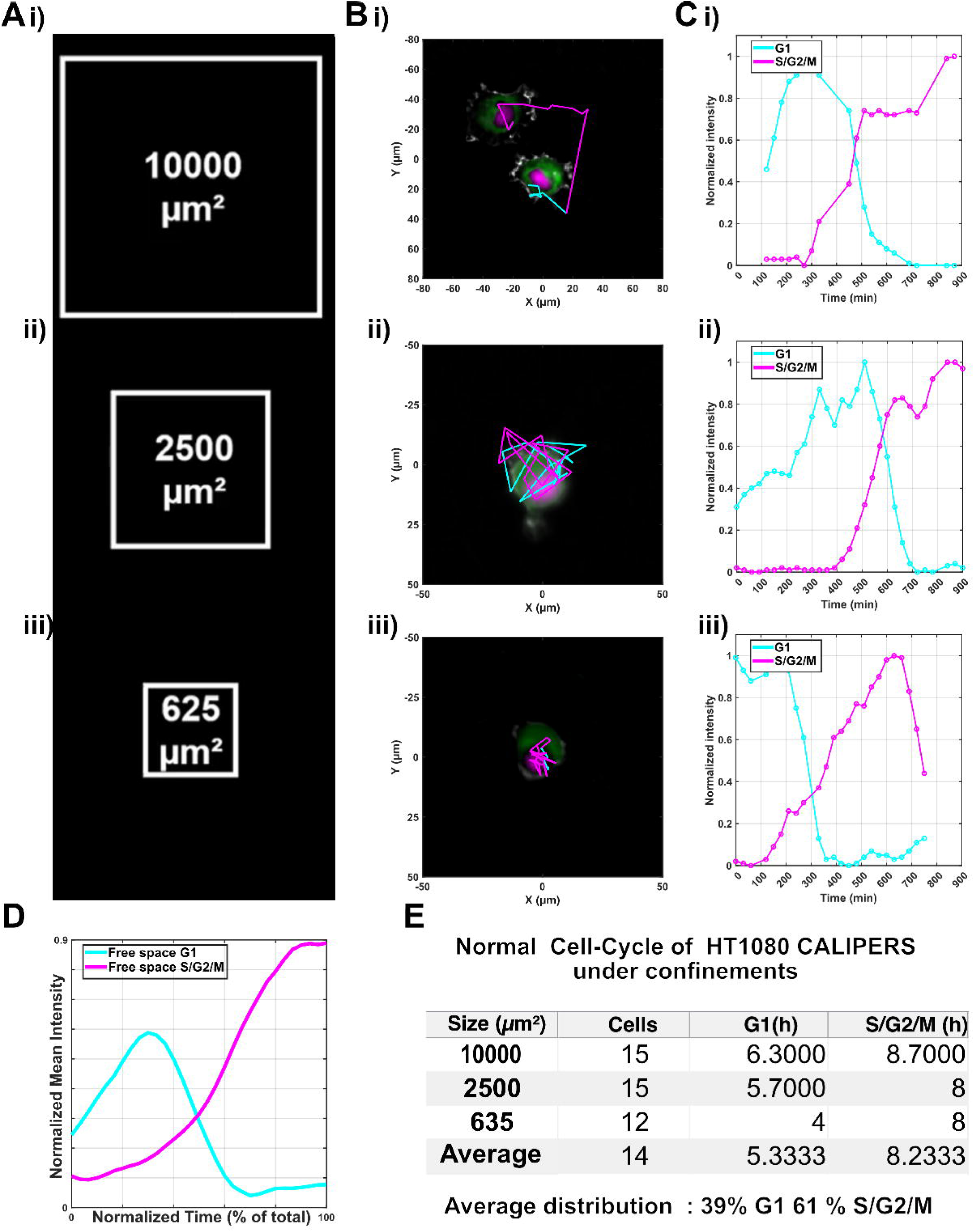

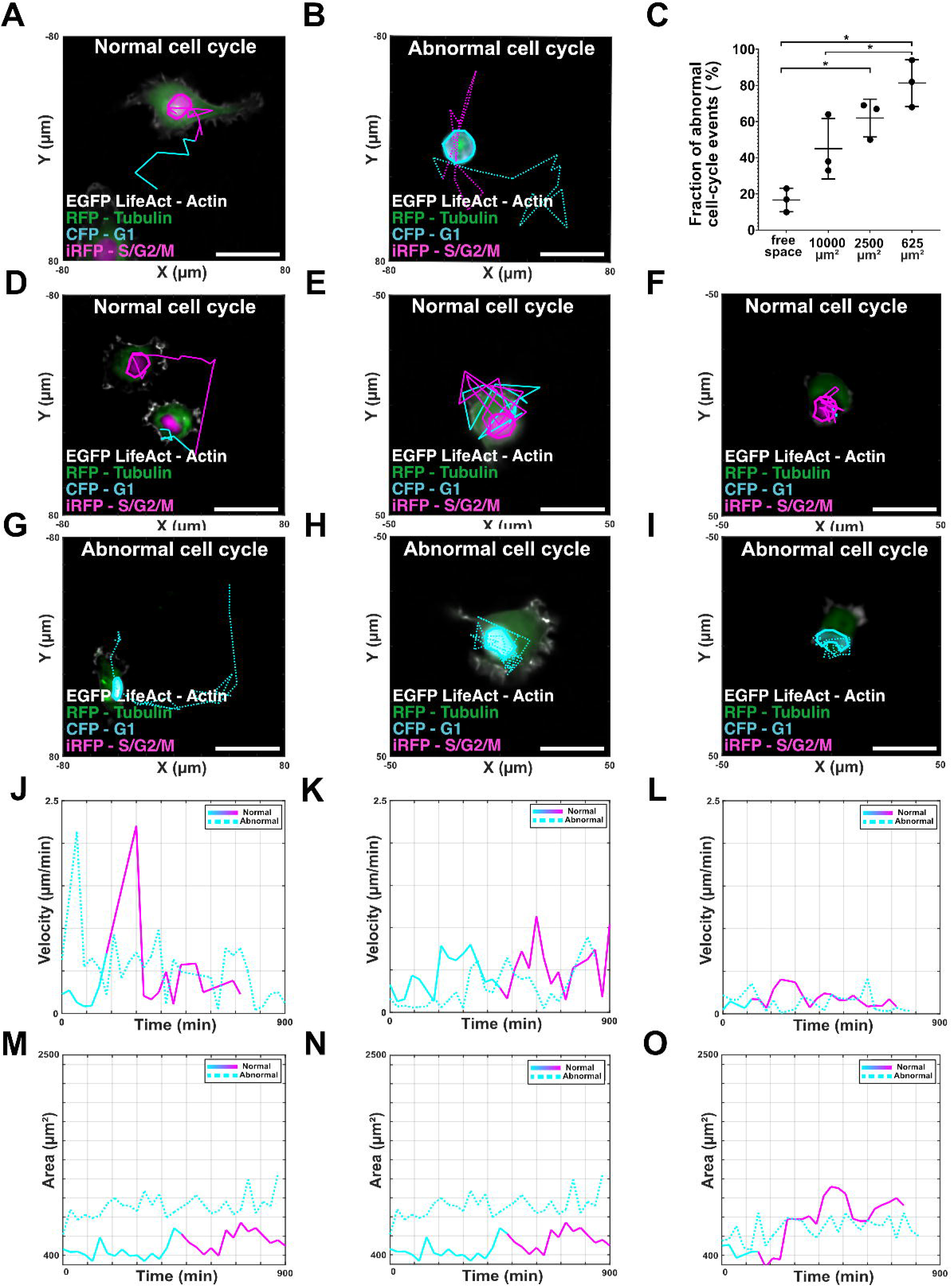

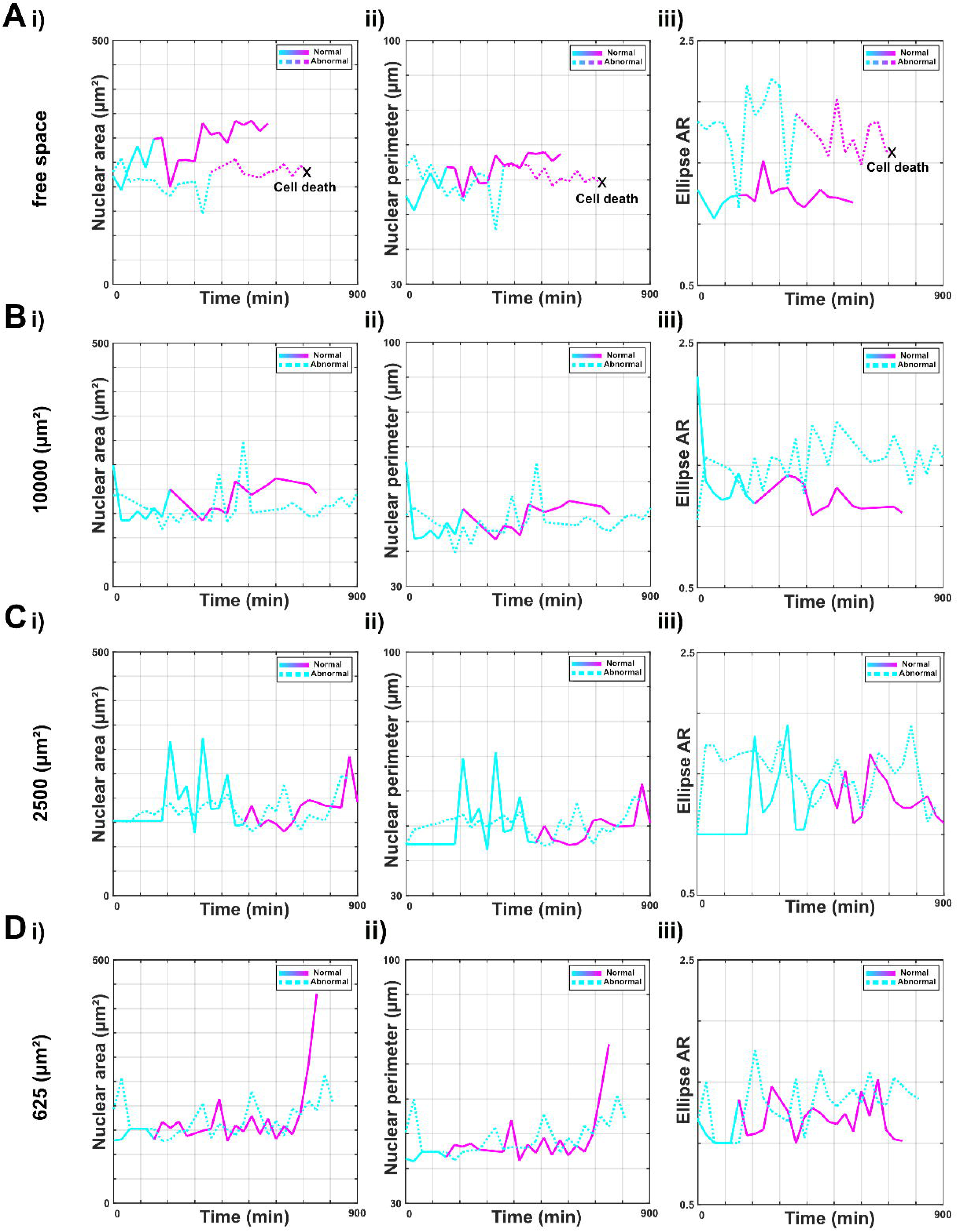

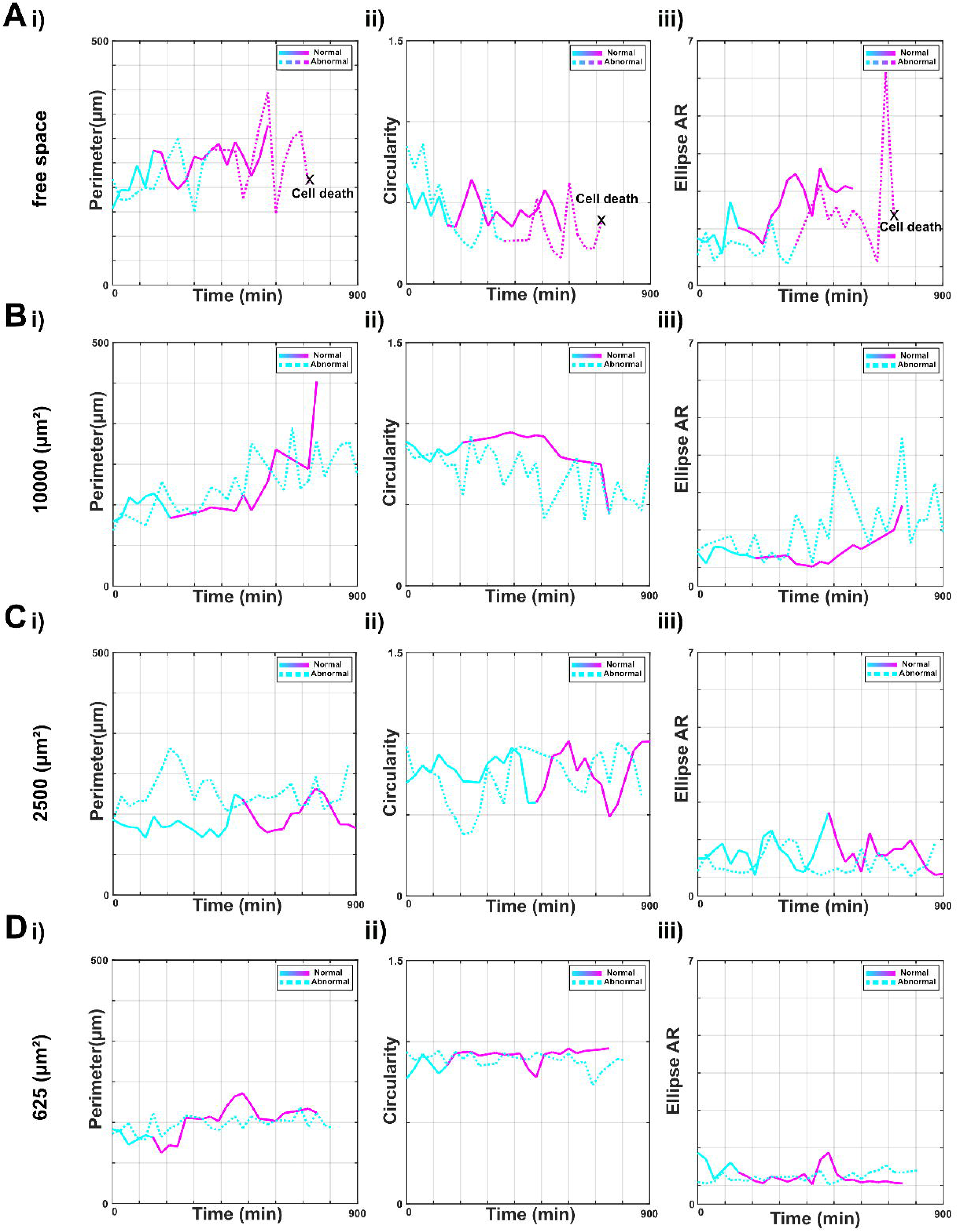

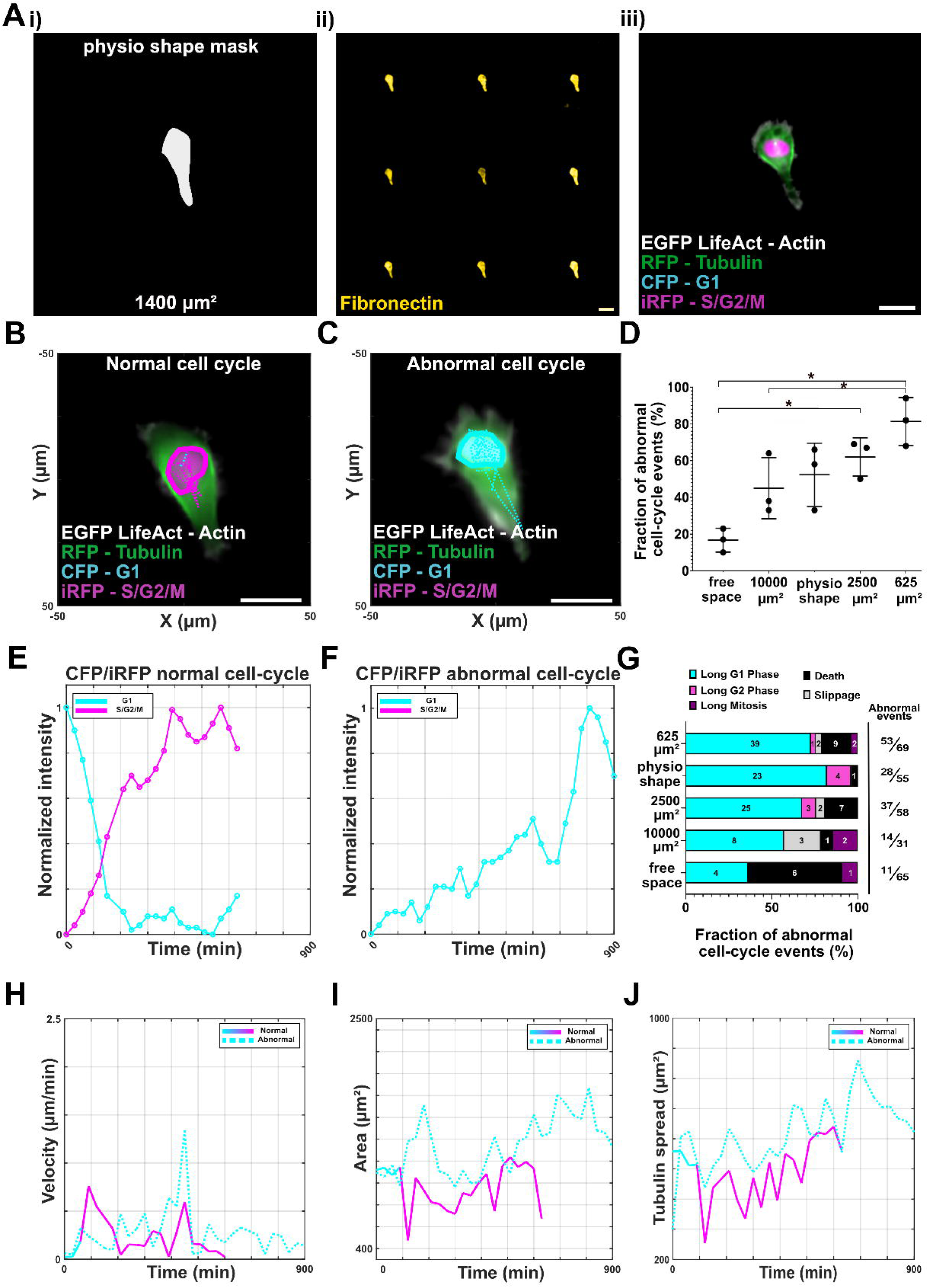

