## Supplementary for "Vertically Integrated System for Tracking and Assessing cell-cycle aware phenotypes under confinement"

### Supplementary figures

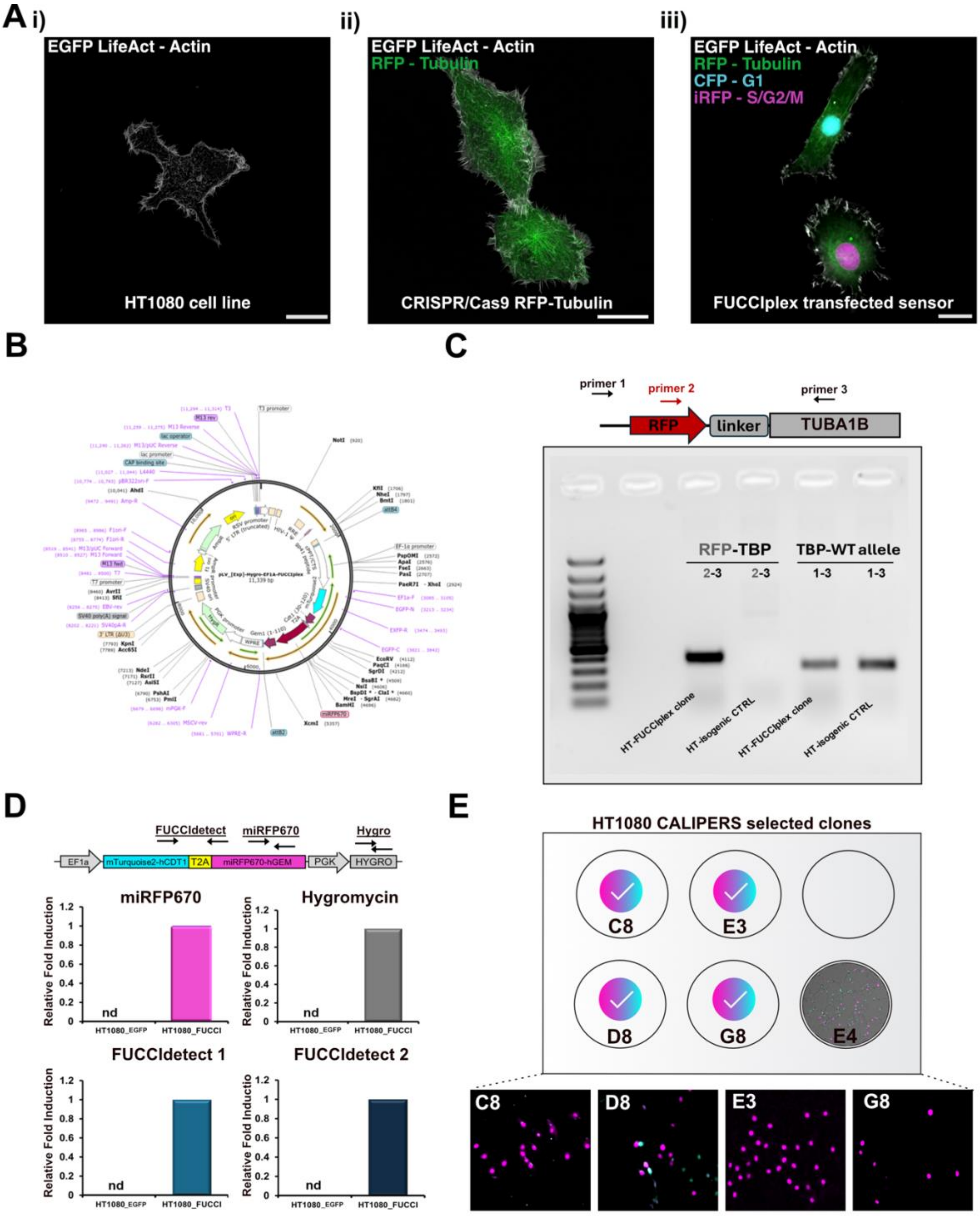

**Figure S1. HT1080 CALIPERS four fluorophores generation**

(A) Representative fluorescence micrographs showing (i) HT1080 cells expressing EGFP LifeAct to visualize actin cytoskeleton, (ii) CRISPR/Cas9-mediated knock-in of RFP-tubulin, and (iii) multiplexed FUCCIplex reporters integrated with the RFP-tubulin line to monitor cytoskeletal structure and cell-cycle phase (CFP for G1, iRFP for S/G2/M). Scale bars, 25 µm. (B) Schematic plasmid map of FUCCIplex construct highlighting insertion sites, promoters, and selection cassettes. (C) Genomic PCR shows successful integration of RFP at the TUBA1B locus in HT1080 CALIPERS clones (primers 2–3), with HT1080 EGFP LifeAct (WT) alleles detected using primers 1–3. CTRL = non-edited HT1080 DNA (HT1080_EGFP). (D) RT-qPCR detects miRFP670, Hygromycin, and FUCCIplex-specific transcripts in HT1080 CALIPERS clone E4 but not in HT1080 EGFP LifeAct cells ( nd = not detected). (E) Four additional stable reference lines for HT1080 CALIPERS: C8, D8, E3, G8, representative imaging under expansion condition within the selected clone E4, as the HT1080 CALIPERS line used for downstream experiments. Scale bars, 100 µm.

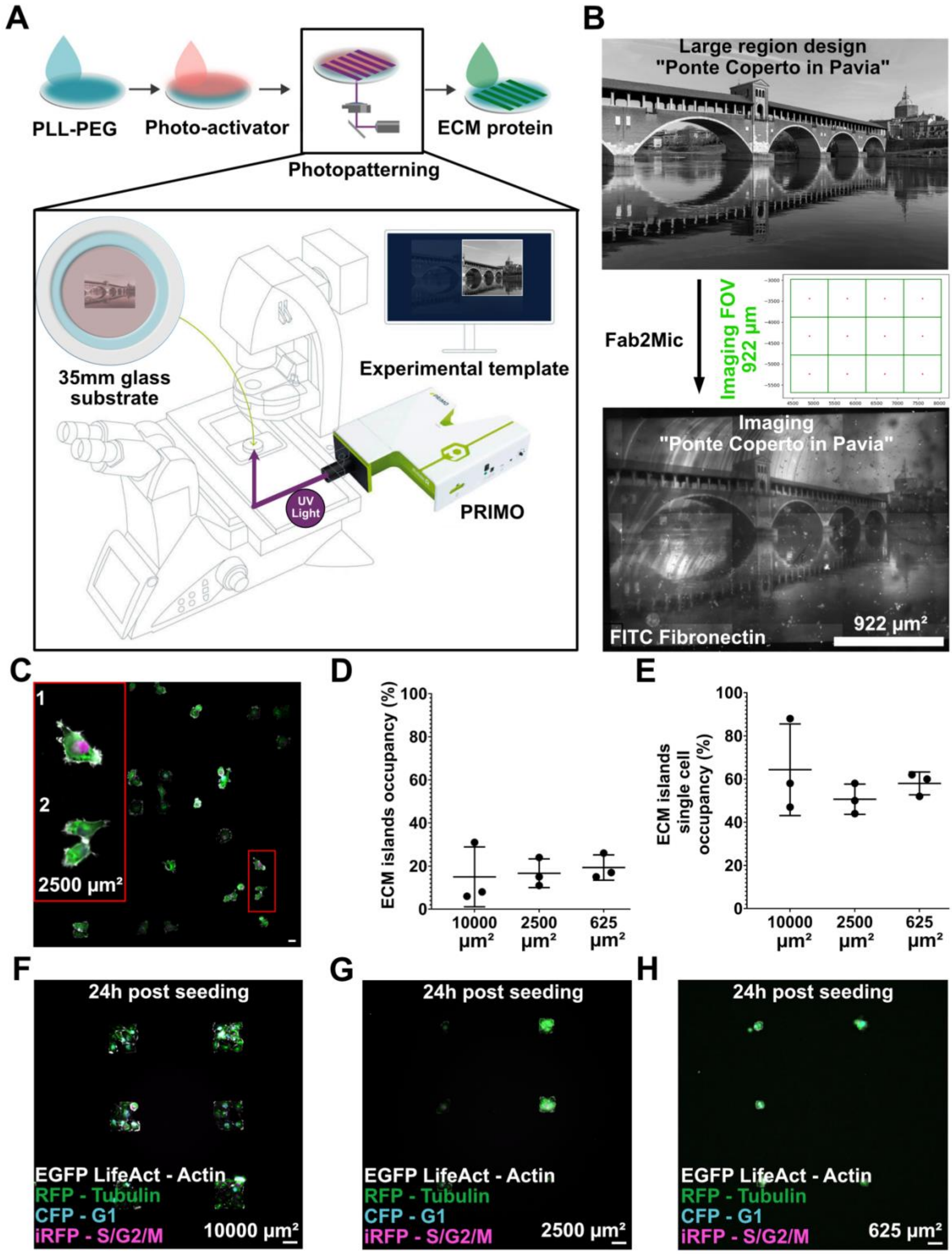
A
PLL-PEG
Photo-activator
Photopatterning
ECM protein
35mm glass substrate
Experimental template
UV Light
PRIMO
B
Large region design
"Ponte Coperto in Pavia"
Fab2Mic
Imaging FOV 922 µm
Imaging
"Ponte Coperto in Pavia"
FITC Fibronectin
922 µm²
C
1
2
2500 µm²
D
ECM islands occupancy (%)
10000 µm²
2500 µm²
625 µm²
E
ECM islands single cell occupancy (%)
F
24h post seeding
EGFP LifeAct - Actin
RFP - Tubulin
CFP - G1
iRFP - S/G2/M
10000 µm²
G
2500 µm²
H
625 µm²

**Figure S2. Fab2Mic validation and single-cell occupancy evaluation in square ECM islands** (A) Schematic of the photopatterning protocol. Glass substrates were coated with PLL-PEG, exposed to photoactivator through the PRIMO UV system, and functionalized with ECM protein. Additionally, the developed Fab2Mic routine aligned fabrication templates with imaging fields of view (FOVs). (B) Example of a large-region design patterned and coated with FITC-fibronectin, representing the "Ponte Coperto in Pavia." The Fab2Mic algorithm generated imaging coordinates (922 μm FOV) for tiled acquisition (See Movie S1). (C) Representative fluorescence images of patterned engineered ECM islands (2500 $\mu m^2$). Insets highlight poor single-cell occupancy and clusters adhering to individual islands. Scale bar, 25 μm. (D) Quantification of the overall engineered ECM island occupancy percentage across designs (10000, 2500, 625 $\mu m^2$) within the defined seeding strategy (E). Single-cell occupancy percentage per island. Data show mean ± SD. (F–H) Representative live-cell imaging of HT1080 CALIPERS cells seeded on ECM islands of (F) 10000 $\mu m^2$, (G) 2500 $\mu m^2$, and (H) 625 $\mu m^2$. Actin (EGFP LifeAct), microtubules (RFP-Tubulin), and cell-cycle reporters (CFP-G1, iRFP-S/G2/M) are shown at 24 hours post-seeding, highlighting overcrowded environments: scale bars, 25 μm.

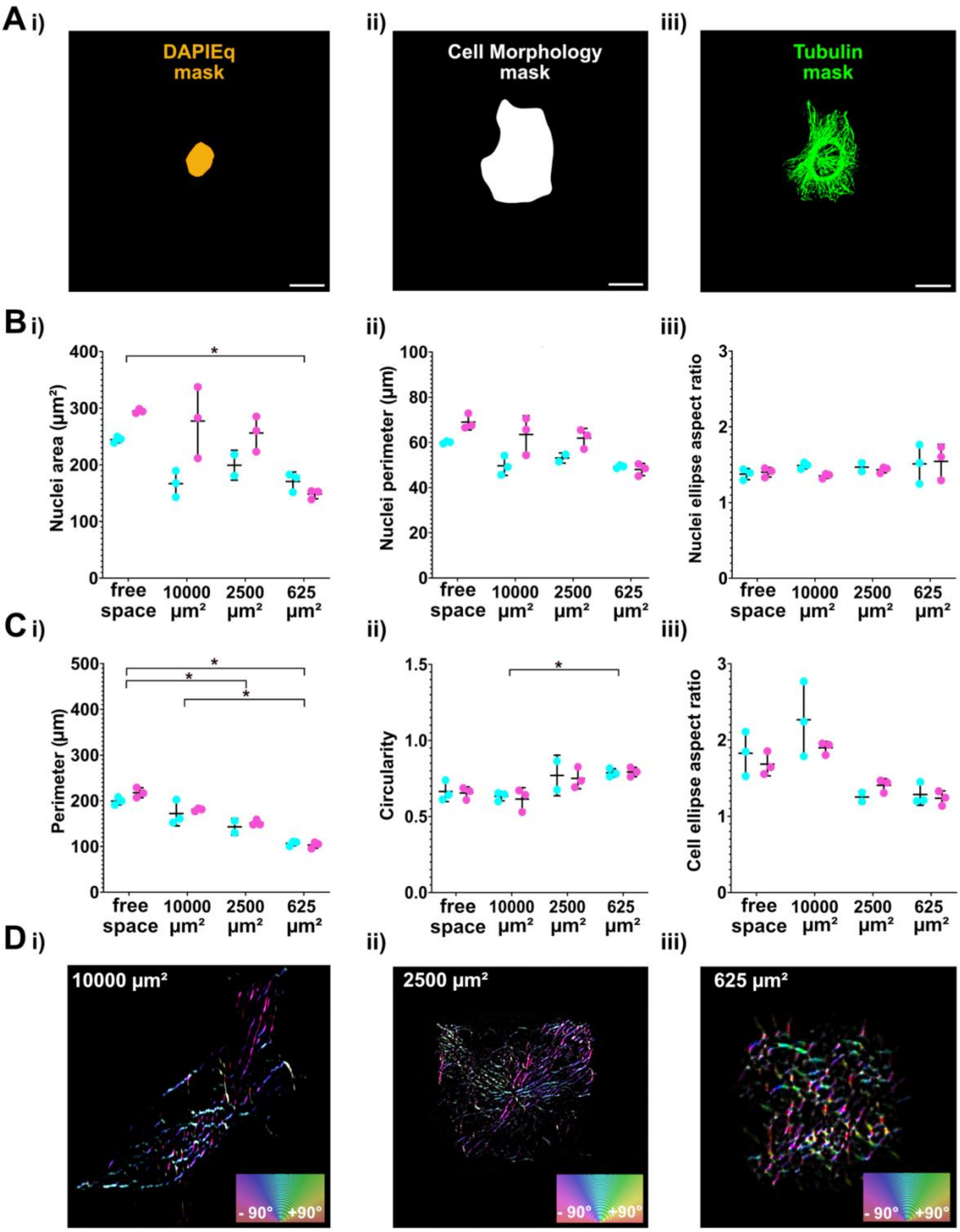


**Figure S3. Additional nuclear and cytoskeletal morphology features across ECM confinements.** (A) Representative segmentation masks used for quantitative analysis: (i) nuclear mask (DAPIEq), (ii) whole-cell morphology mask, and (iii) tubulin cytoskeleton mask. Scale bars, 25 µm. (B) Nuclear morphology static metrics across free space and confined geometries (10000, 2500, 625 µm² islands) in three independent experiments: (i) nuclear area, (ii) nuclear perimeter, and (iii) nuclear ellipse aspect ratio. Data show mean ± SD; (*$p < 0.05$). (C) Whole-cell morphological descriptors across conditions: (i) cell perimeter, (ii) circularity, and (iii) ellipse aspect ratio. Data show mean ± SD; (*$p < 0.05$). (D) Orientation maps[73] of tubulin filaments in cells confined on ECM islands of (i) 10000, (ii) 2500, and

(iii) 625 µm². Orientation vectors are color-coded according to the indicated scale bars (–90° to +90°). Insets show color scales.

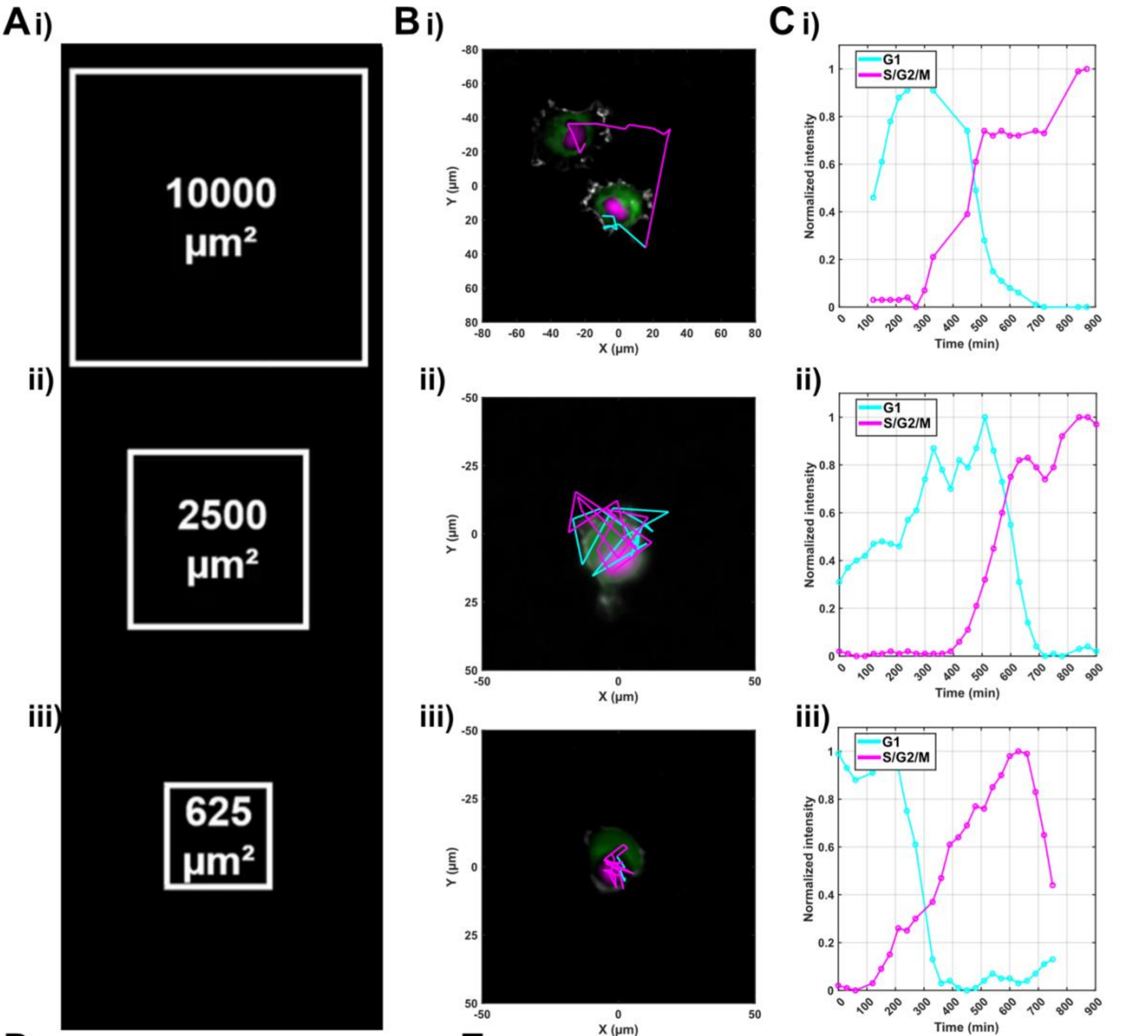


Normal Cell-Cycle of HT1080 CALIPERS under confinements

| Size (µm²) | Cells | G1(h) | S/G2/M (h) |
|---|---|---|---|
| 10000 | 15 | 6.3000 | 8.7000 |
| 2500 | 15 | 5.7000 | 8 |
| 635 | 12 | 4 | 8 |
| Average | 14 | 5.3333 | 8.2333 |

Average distribution : 39% G1 61 % S/G2/M

**Figure S4. Normal cell-cycle dynamics of HT1080 CALIPERS cells under ECM confinement.** (A) Representative ECM confinement geometries of 10000 µm² (i), 2500 µm² (ii), and 625 µm² (iii) patterned islands used for analysis. (B) Trajectories of representative HT1080 CALIPERS cells undergoing normal cell-cycle in each confinement, showing actin (EGFP LifeAct), tubulin (RFP), and FUCCIplex reporters (CFP for G1, iRFP for S/G2/M). Migration paths are overlaid. (C) Time-resolved FUCCIplex intensity profiles (CFP = cyan, iRFP = magenta) corresponding to the cells shown in (B), with sequential progression through G1 and S/G2/M phases. (D) Average normalized FUCCIplex intensity traces of HT1080 CALIPERS cells in free space, showing mean temporal dynamics of G1 and S/G2/M phases. (E) Summary table of normal cell-cycle duration under confinement, reporting average times spent in G1 and S/G2/M phases across 10000, 2500, and 625 $\mu m^2$ ECM-islands. The overall distribution across all conditions was 39% G1 and 61% S/G2/M.

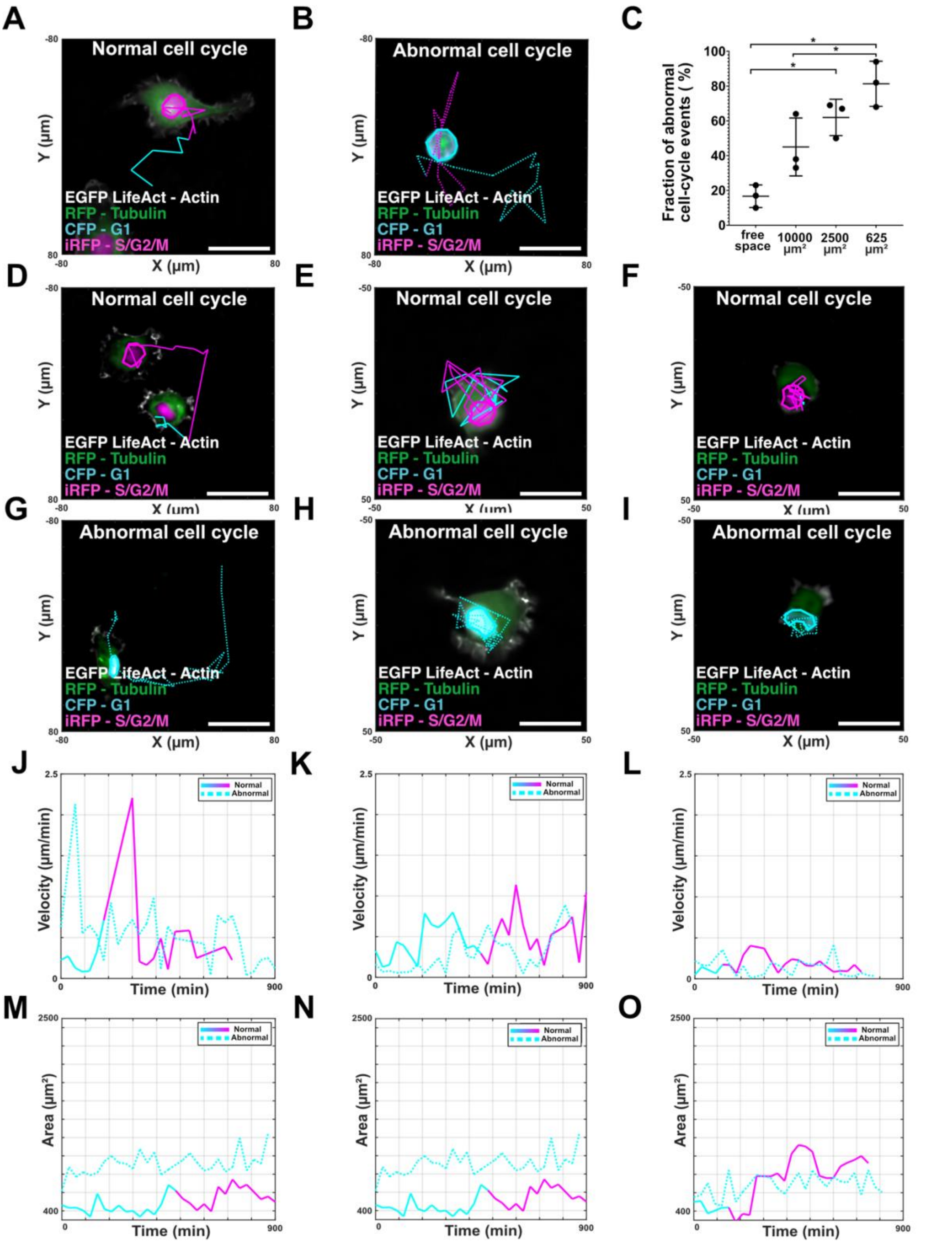


**Figure S5. Extended phenotyping of normal vs. abnormal cell-cycle events under ECM confinement.** (A–B) Representative trajectories of CALIPERS cells in free space, undergoing (A) normal or (B) abnormal cell-cycle events. See Movie S2 (D, G, J, M) Representative trajectories and quantitative dynamics in 10000 µm² confinement, showing (D) normal, (G) abnormal cell-cycles, with corresponding (J) velocity and (M) area dynamics. (E, H, K, N) Representative trajectories and dynamics in a 2500 µm² confinement, showing (E) normal and (H) abnormal cycles, with (K) velocity and (N) area profiles. (F, I, L, O) Representative trajectories and dynamics in 625 µm² confinement, showing (F) normal, (I) abnormal cycles, with (L) velocity and (O) area profiles. Cells express EGFP

LifeAct (actin), RFP-tubulin, CFP (G1), and iRFP (S/G2/M). Migration trajectories are overlaid. Scale bars, 25 μm. (C) Quantification of abnormal cell-cycle events across confinement conditions. Data show mean ± SD; *$p < 0.05$.

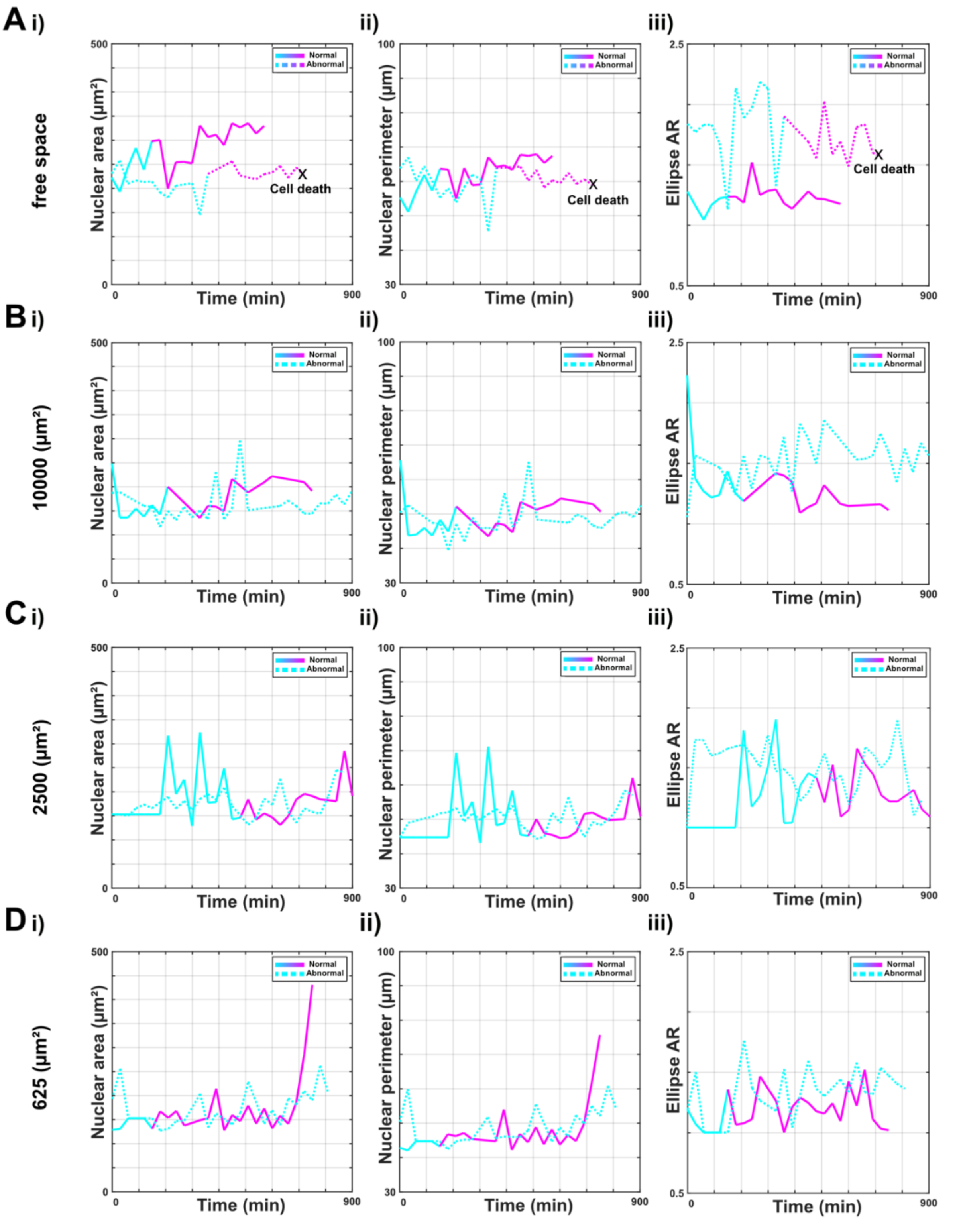


**Figure S6. Nuclear dynamics across different square ECM sizes in normal vs. abnormal cell-cycles.**

(A–D) Time-resolved nuclear morphology metrics of CALIPERS cells tracked over 15 h in free space (A), 10000 μm² islands (B), 2500 μm² islands (C), and 625 μm² islands (D). For each condition, nuclear (i) area, (ii) perimeter, and (iii) ellipse aspect ratio (AR) are plotted for cells undergoing normal (solid line) and abnormal (dashed line) cell-cycle. Cell death is indicated where applicable (X).

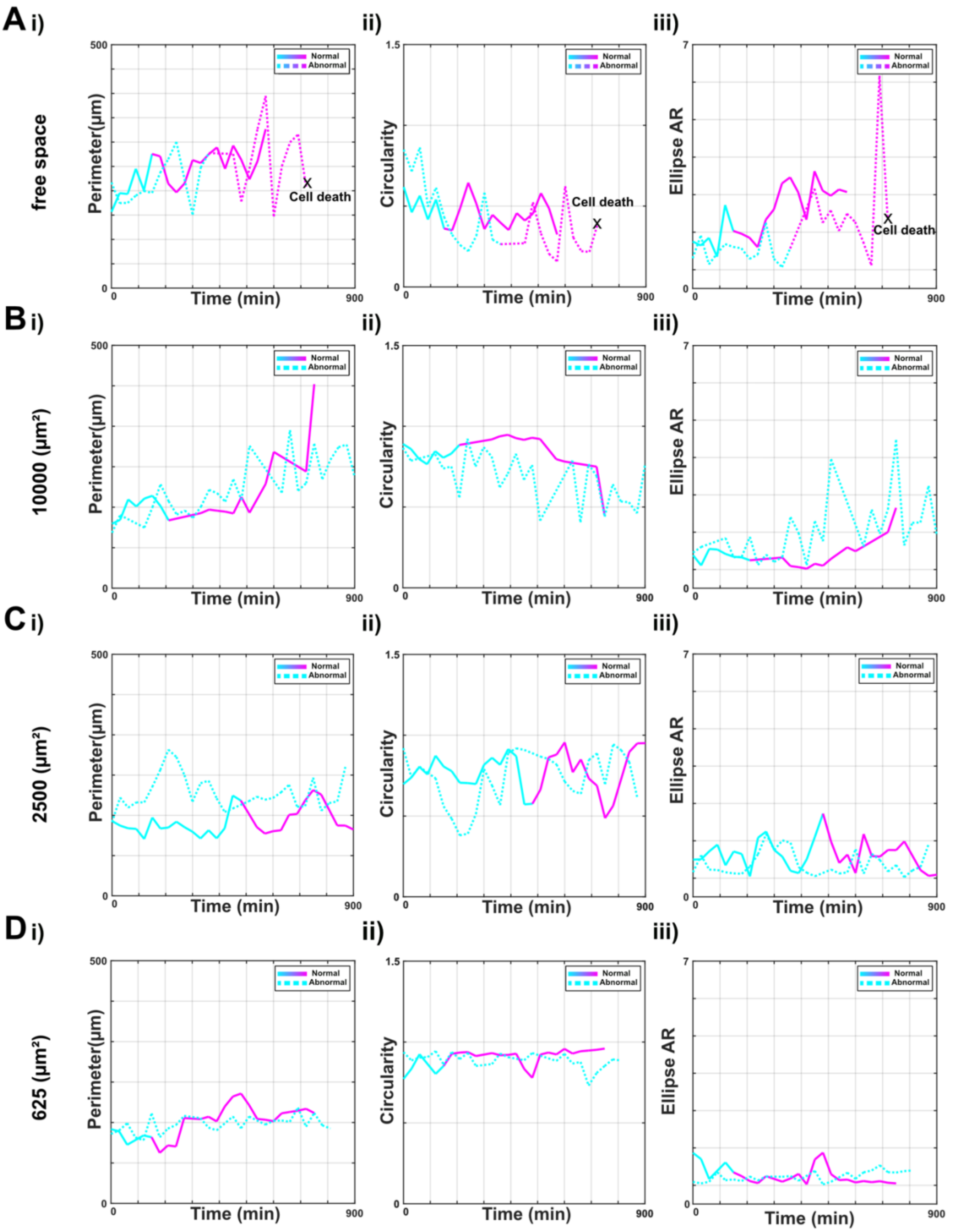


**Figure S7. Cell morphological dynamics across different square ECM sizes in normal versus abnormal cell-cycles.**

(A–D) Time-resolved morphological metrics of HT1080 CALIPERS cells tracked over 15 h in free space (A), 10000 μm² (B), 2500 μm² (C), and 625 μm² (D) ECM islands. For each condition, (i) cell perimeter, (ii) circularity, and (iii) ellipse aspect ratio (AR) are plotted for cells undergoing normal (solid line) and abnormal (dashed line) cycles. Instances of cell death are indicated where applicable (X).

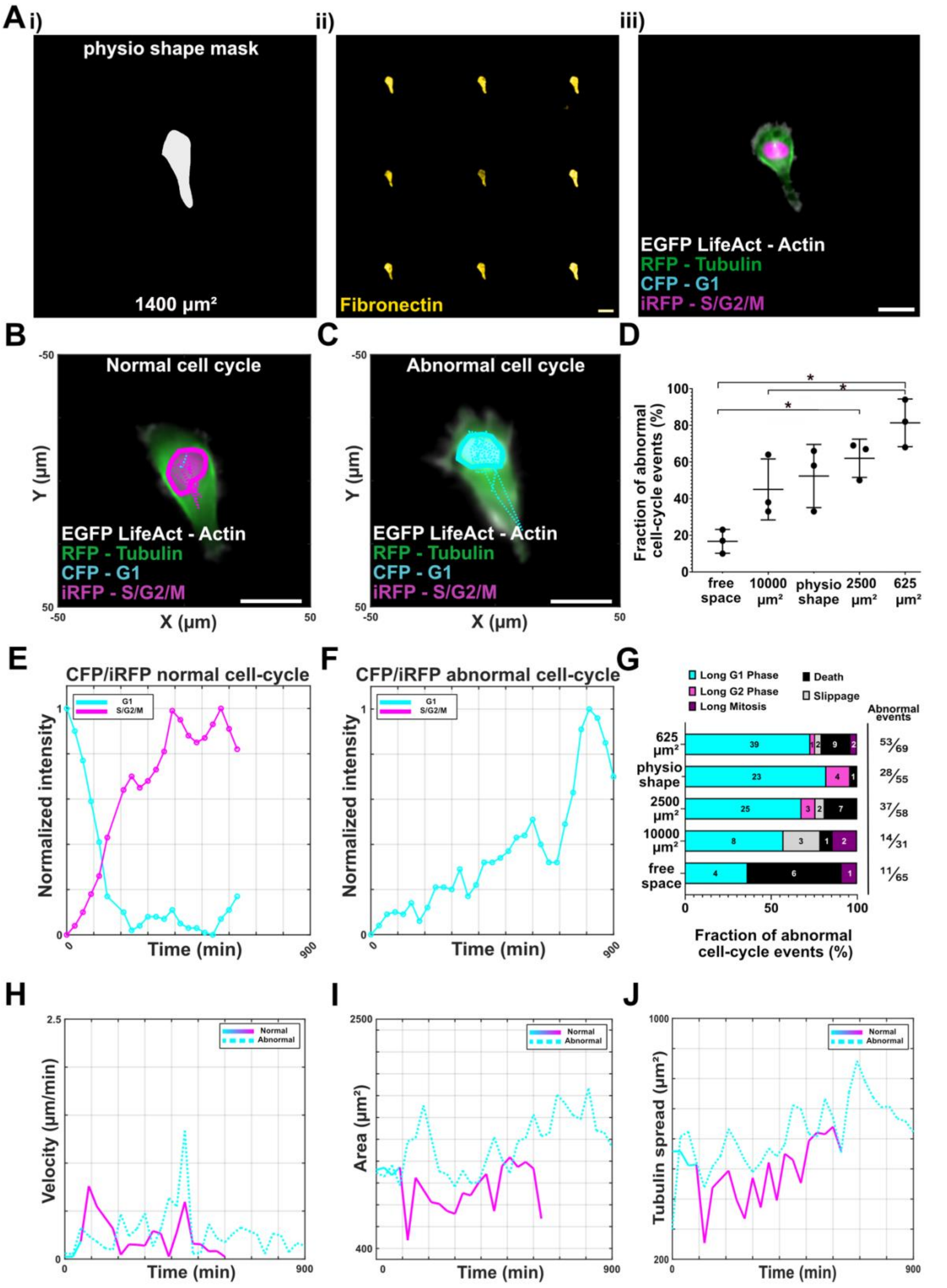


**Figure S8. Physiologically inspired ECM island and related cell-cycle abnormalities.** (A) Representative masks and fluorescent validation of HT1080 CALIPERS cells on physio-shaped ECM islands (Physiological mean area size, 1400 µm²). (i) Design of a physio-shaped island, (ii) Photopatterning validation with a FITC-fibronectin-coated array of physio shape islands, and (iii) single cell expressing EGFP LifeAct (actin), RFP-tubulin, CFP (G1), and iRFP (S/G2/M) adhered to physio-shaped engineered ECM island. Scale bars, 25 µm. (B–C) Example cells undergoing (B) normal and

(C) abnormal cell-cycle events under confinement. Scale bars, 25 µm. (D) Quantification of abnormal cell-cycle events across free space, 10000 µm², physio-shaped, 2500 µm², and 625 µm² islands. Data show mean ± SD; *$p < 0.05$. (E–F) Representative CFP/iRFP intensity profiles from cells undergoing (E) normal or (F) abnormal cycles. (G) Distribution of abnormal cell-cycle events across different confinement geometries, defined as Long G1, Long S/G2/M, long mitosis, S/G2/M-G1 slippage, or cell death. (H–J) Quantitative comparison of motility and morphology dynamics between normal and abnormal cycles: (H) velocity, (I) cell area, and (J) tubulin spread over time under physio-shape confinement.

### Supplementary movies

**Movie S1:** Correlative fabrication-to-microscopy pipeline

**Movie S2:** Normal vs. abnormal cell-cycle events in free space and square ECM confinements
